# Planar cell polarity-directed cell crawling drives polarized hair follicle morphogenesis

**DOI:** 10.1101/2025.11.13.688366

**Authors:** Rishabh Sharan, XinXin Du, Liliya Leybova, Anyoko Sewavi, Abhishek Biswas, Danelle Devenport

## Abstract

During epithelial morphogenesis, cell polarity aligns individual cell behaviors into collective motions that shape developing tissues. Here, we combine experiments with computational modeling to investigate how cell-scale forces oriented by Planar Cell Polarity (PCP) direct the collective, counter-rotational cell flows that occur during hair placode morphogenesis. We rule out that PCP directs apical neighbor exchanges, as junctional myosin and PCP protein localization are not co-correlated with junction shrinkage. Instead, we find that PCP directs anterior-directed crawling of placode cells along the basal surface of the tissue through a mechanism that requires cell crawling regulator Rac1. Modeling the placode as a continuum viscoelastic fluid, we find that active forces from cell crawling at the basal surface is sufficient to generate the experimentally observed counter-rotational cell motion at the apical surface. Our results show an unexpected role for PCP in epithelial morphogenesis, centering the basal surface as the site of force generation.

## Introduction

Understanding how cells coordinate their individual behaviors to generate tissue-scale morphogenetic change is a central question in developmental biology. Active forces from cell behaviors such as shape changes, proliferation, migration, and intercalation are sources of stress that ultimately deform tissue sheets, creating bends, folds, or elongations that contribute to an organ’s final shape. In epithelia, which are polarized along both apical-basal and planar axes of a tissue, cell shape changes and movements often arise through the asymmetric localization of force-generating cytoskeletal networks, whose position within cells determines the magnitude and directionality of epithelial deformations. For example, apical positioning of contractile actomyosin networks constricts cells apically resulting in a concave tissue bend, or invagination (Ettensohn, 1985). Additionally, localization of actin polymerization machinery to one pole of the cell in the plane of the tissue can promote directed migration or tissue extension (Sanchez-Corrales et al., 2018; Williams et al., 2022; Zallen and Wieschaus, 2004). Thus, to ensure an epithelium bends or elongates reproducibly, individual cells must polarize the force generating machinery to the correct subcellular location and coordinate this behavior across the cell collective.

In classic models of epithelial tissue elongation, such as *Drosophila* germband extension and amniote neurulation, planar polarity systems localize actomyosin cables to anterior-posterior cell junctions, promoting cell intercalation and tissue convergence to elongate the embryonic body axis (Blankenship et al., 2006; Butler and Wallingford, 2018; Nishimura et al., 2012; Williams et al., 2014; Zallen and Wieschaus, 2004). Actomyosin cables are positioned at the apico-lateral edges of intercalating cells, suggesting that forces at the apical surface drive intercalation. However, events at the basal surface have also been implicated in force generation (Sun et al., 2017; Williams et al., 2014; Williams-Masson et al., 1998). There, intercalating cells display crawling behaviors on their basal sides, and neighbor exchanges often initiate basally, rather than apically. These studies challenged the classic model for planar polarity-driven morphogenesis, centering the basal, rather than apical surface as the site of tissue-scale force generation.

Mammalian hair follicles serve as useful miniorgans for deciphering how cell polarity regulates force generation during morphogenesis. Hair follicles begin as circular epithelial thickenings known as hair placodes, which consist of at least two distinct cell types, inner and outer cells, organized as a radially symmetric disc. During placode budding, stereotyped in-plane cell rearrangements reshape the placode into an epithelial bud that bends with an anterior-facing tilt (Fig. 1A). We term this process *hair placode polarization*, and it results in the characteristic, uniform alignment of body hairs across the epidermal surface (Cetera et al., 2018; Devenport and Fuchs, 2008). Planar cell polarity (PCP) directs the in-plane, counter-rotational cell rearrangements that reposition inner cells toward the anterior end of the bud and sweep outer cells posteriorly (Fig. 1A). In the absence of PCP function, placode cell rearrangements are reduced or randomized, resulting in hair follicles that are misaligned relative to the body axes (Cetera et al., 2018; Devenport and Fuchs, 2008; Wang et al., 2010). While it is clear that PCP is required for the stereotyped pattern of cell motion, how PCP directs the in-plane intracellular forces that drive hair placode polarization, and whether these occur apically or basally, remains unknown.

**Figure 1.**
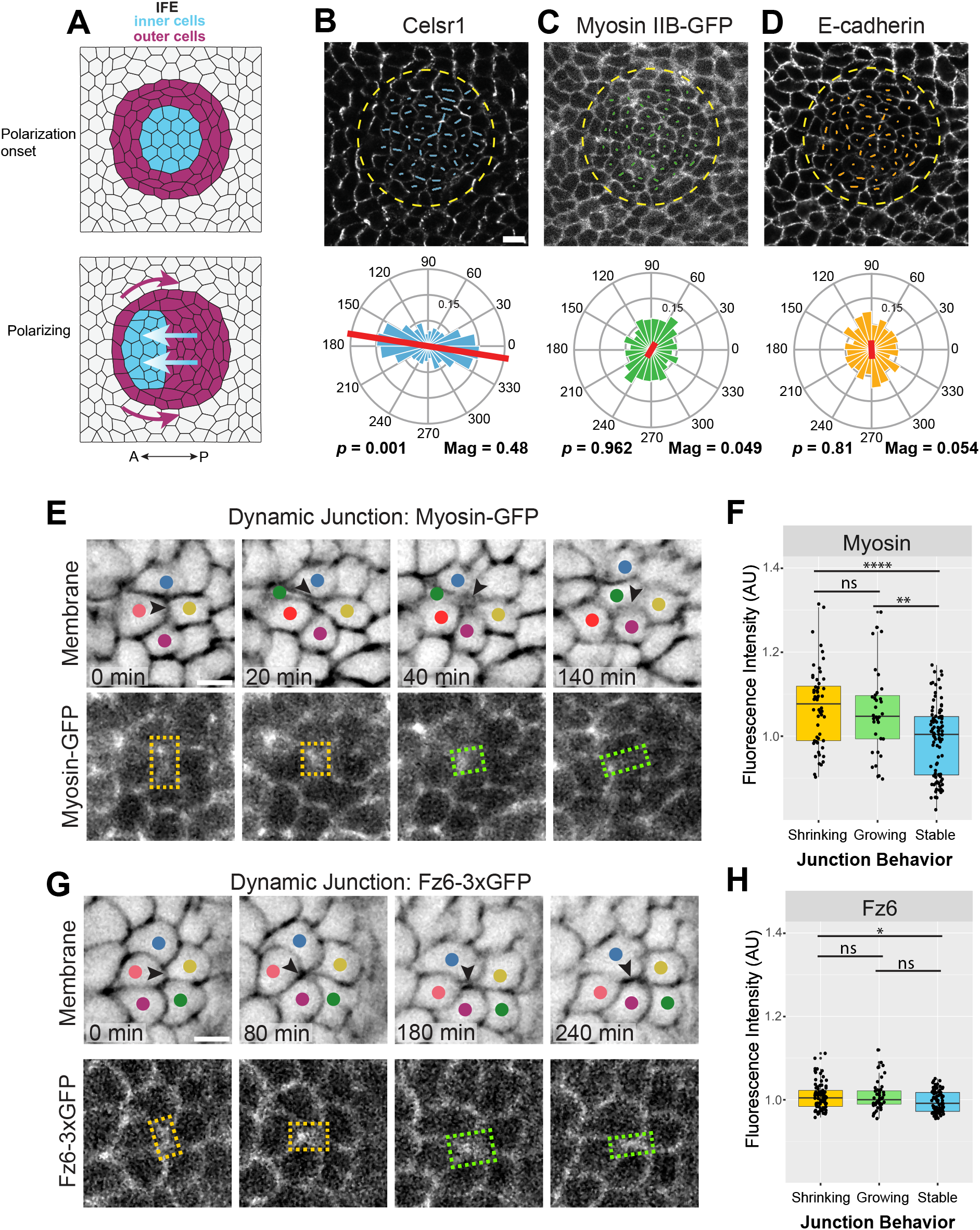
Spatial and temporal distributions of Myosin and PCP components do not correlate with junction remodeling during hair placode polarization. A) Schematic depicting the hair placode at the onset of polarization (top) and during polarization (bottom). Planar view is shown. Inner cells (blue) move towards the anterior whereas outer cells (magenta) are displaced toward the posterior. B-D) Representative planar views of hair placode from *E*15.5 embryo expressing Myosin IIB fluorescently-tagged with GFP. Skin explants were immunolabeled with antibodies against Celsr1 and E-cadherin. Images are overlaid with colored lines representing the axis and magnitude of nematic polarity for each cell by line orientation and length (top panels). Quantification of polarity distributions are displayed below as circular histograms (bottom panels). Direction of red lines corresponds to the average angle of polarity, and length of red lines corresponds to the average magnitude of polarity, whose value is given below (Mag). Permutation test by random resampling of data performed to obtain *p*-value. Anterior to the left. n = 465 cells, N = 6 hair placodes. Scale bar: 10 µm. E) Representative time-lapse images from *E*15.5 skin explants co-expressing membrane-tdTomato (top panel) and Myosin IIB-GFP (bottom panel). Images are taken from the center of a developing hair placode during polarization (see Fig. S1A). In (E) a dynamic junction (arrowhead) between cells undergoing a rearrangement is shown (shrinking – orange boxes, growing – green boxes). Scale bar: 5 µm. G) Representative time-lapse images from *E*15.5 skin explants co-expressing membrane-tdTomato (top panel) and Fz6-3xGFP (bottom panel). Images are taken from the center of a developing hair placode during polarization. In (F), a dynamic junction (arrowhead) between cells undergoing a rearrangement is shown (shrinking – orange boxes, growing – green boxes). Scale bar: 5 µm. F and H) Fluorescence intensities of Myosin IIB-GFP (F) or Fz6-3xGFP (H) at stable and dynamic junctions, where dynamic junction time points are categorized as shrinking or growing based on junction length difference with previous timepoint. Fluorescence intensity values are normalized to the mean of all stable junctions across all time points per time-lapse image. n=9 junctions (stable and dynamic), N = 3 hair placodes.

Given the junctional localization of PCP proteins in cells of the skin epidermis, we initially hypothesized that PCP directs actomyosin to apico-lateral cell boundaries in hair placodes to promote junction remodeling and in-plane cell rearrangements. However, through fixed and live-imaging of myosin during placode polarization, we show that myosin is not asymmetrically localized to PCP-enriched junctions. Additionally, we find that although myosin enrichment correlates temporally with junction shrinkage in intercalating cells, PCP protein localization does not. Instead, through 3D volumetric reconstructions and live imaging, we find that central, inner cells of the hair placode display oriented protrusions along the basal surface, suggesting that cell crawling may, in fact, be the driving force of placode polarization. Through continuum mechanics modeling we show that cell crawling is sufficient to drive the experimentally observed counter-rotational pattern of cell movements during hair placode polarization. Moreover, we show that non-uniform material properties within the placode and surrounding tissue allow for stronger and more robust formation of apical counter-rotational flows. Using inhibitors targeting the cell crawling regulator Rac1, we show that cell crawling is required, indeed, for hair placode polarization. We conclude that the function of epidermal PCP proteins is to direct cell crawling of placode cells toward the anterior, resulting in the collective cell rearrangements that accompany hair placode polarization.

## Results

### Myosin does not correlate with PCP protein localization at cell junctions during hair placode polarization

In many systems, cell intercalation in epithelial tissues is driven by polarized actomyosin-based junction contraction (Bertet et al., 2004; Blankenship et al., 2006; Butler and Wallingford, 2018; Nishimura et al., 2012; Sanchez-Corrales et al., 2018; Williams et al., 2014; Zallen and Wieschaus, 2004). Therefore, we initially hypothesized that PCP proteins may promote neighbor exchange in polarizing hair placodes by directing myosin contractility to PCP-enriched junctions. If our hypothesis is correct, myosin is expected to exhibit polarized localization or activity at PCP-enriched anterior-posterior junctions (also referred to as vertical junctions), as observed in chick and *Xenopus* neural tube closure (Butler and Wallingford, 2018; Nishimura et al., 2012). Furthermore, we would expect PCP protein localization to correlate with both myosin localization and junction shrinkage.

To visualize myosin localization, we utilized a previously generated transgenic mouse strain in which GFP is endogenously fused to the N-terminus of non-muscle myosin heavy chain II-B (Myosin IIB-GFP) (Bao et al., 2007). Whole mount epidermal explants from Myosin IIB-GFP expressing embryos were immunolabeled with antibodies against Celsr1 to visualize PCP protein localization (Devenport and Fuchs, 2008), and against E-cadherin to mark epithelial junctions. Following semi-automated segmentation of cell edges (Pachitariu and Stringer, 2022), we quantified the orientation and magnitude of Celsr1, Myosin IIB-GFP, and E-cadherin polarization within the epithelial plane using TissueAnalyzer (Aigouy et al., 2010; Aigouy et al., 2016; Aw et al., 2016). As expected, Celsr1 displayed asymmetric localization and was enriched along vertical junctions in placode cells (Fig. 1B), as has been shown previously (Devenport and Fuchs, 2008). This is in contrast with E-cadherin that is uniformly distributed around epithelial cell membranes (Fig. 1D) (Devenport and Fuchs, 2008). Surprisingly, we found that Myosin IIB-GFP was not asymmetrically distributed to vertical junctions and was uniformly distributed around the cell membrane (Fig. 1C). We additionally tested whether activated myosin II was polarized by examining the localization of phosphorylated myosin light chain (p-MLC). We found that p-MLC mainly accumulated at tricellular vertices in the placode epithelium. Its distribution was not planar polarized, nor did it co-localize with Celsr1 (Fig. S1). Thus, unlike Celsr1 and the other core PCP proteins (Devenport and Fuchs, 2008; Basta et al., 2021), non-muscle myosin II is not planar polarized at vertical junctions of the placode epithelium.

We then considered the possibility that PCP proteins and myosin dynamics might correlate specifically with junction remodeling, as previously observed during *Xenopus* neural tube closure (Butler and Wallingford, 2018). To test whether PCP proteins and myosin dynamics correlate with junction shrinkage, we generated mouse embryos expressing Myosin IIB-GFP or Fz6-3xGFP together with membrane-tdTomato (mTomato) (Bao et al., 2007; Basta et al., 2021; Muzumdar et al., 2007), which enabled us to simultaneously visualize cell edges together with myosin or a PCP protein as hair placode cells undergo neighbor exchanges (*Myosin IIB-GFP; mTomato* and *Fz6-3xGFP; mTomato*). Time-lapse imaging was performed on live skin explants from *E*15.5 *Myosin IIB-GFP; mTomato* and *Fz6-3xGFP; mTomato* embryos for approximately 18 hours to capture cell rearrangements that occur during hair placode polarization (Fig.S2A, Movie S1, S2). Using the membrane channel to visualize junction behavior, we categorized junctions as stable or dynamic. Stable junctions were defined as junctions that were not part of a neighbor exchange. Dynamic junctions were defined as those which were part of a neighbor exchange or whose length underwent significant growth or shrinkage. For a given dynamic junction, each timepoint was classified as shrinking or growing based on the junction length difference with the previous timepoint. For each junction and timepoint, we measured the mean Myosin IIB-GFP or Fz6-3xGFP fluorescence intensity in the junction region (Fig. 1E, G and Fig. S2B, C). These values were normalized to the average stable junction intensity for each time-lapse movie. As expected, Myosin IIB-GFP intensity was significantly higher during junction shrinkage and growth compared to stable junctions (Fig.1F). Conversely, Fz6-3xGFP intensity was similar between junction behaviors (Fig. 1H). Although a statistically significant difference was observed between Fz6-3xGFP shrinking and stable junction states, the effect size was notably smaller compared to the difference between Myosin IIB-GFP groups. As Myosin IIB-GFP intensity correlated with junction behavior, but Fz6-3xGFP intensity did not, we conclude that it is unlikely that PCP proteins promote neighbor exchanges in the hair placode by directing myosin localization to epithelial junctions.

### Placode epithelial cells form anterior-directed protrusions from their basal surfaces during placode polarization

Thus far, our studies of cell rearrangements have focused on events occurring at the apico-lateral region of cell junctions mainly because in other systems, forces driving junction contraction are localized apico-laterally (Blankenship et al., 2006; Hashimoto et al., 2015; Lienkamp et al., 2012; Nishimura et al., 2012; Sanchez-Corrales et al., 2018). Considering the possibility that cell movements in the placode could be driven by events positioned elsewhere along the apical-basal axis, we generated 3D reconstructions of polarizing hair placodes (Fig. 2A, Movie S3) (Leybova et al., 2024). In volumetric reconstructions of individual epithelial cells, we found that, surprisingly, cells of polarizing placodes formed pronounced protrusions at their basal surface that are anterior directed, resulting in a slanted epithelial cell shape (Fig. 2B, Movie S4). To measure the direction and magnitude of basal surface displacement relative to the apical surface, we calculated the Basal-Apical Offset, which we define as the difference between the x-component of the centroid for the apical (*x*_*A*_) and basal (*x*_*B*_) surfaces (Δ*x* = *x*_*A*_ − *x*_*B*_). Hence, a positive (negative) value of the Offset corresponds to the basal (apical) surface being more anterior (Fig. 2C). As expected, cells of the non-placode, interfollicular epithelium (IFE) did not show a strong Offset in either direction. In the polarizing placode, nearly all inner cells show an Offset of the basal surface toward the anterior, while outer cells have a broad distribution of Offsets (Fig. 2D). These results for outer cells are consistent with our previous observations that outer cells are distinctly shaped from other cells, with their apical surfaces reaching inward and above inner cells (Leybova et al, 2024). Notably, inner cells of the polarizing placode were the only group with a distribution of Offsets where the mean was outside of one standard deviation from 0. This means that inner cells, and not outer cells, during polarization have a statistically significant anterior displacement of the basal relative to the apical surface (Fig. 2D). These results suggest that inner cells may be crawling along the basal surface towards the anterior. From this, we hypothesize that the counter-rotational pattern of cell motion during placode polarization may be due to the anterior-directed crawling of inner cells that displace the surrounding outer cells towards the posterior.

**Figure 2.**
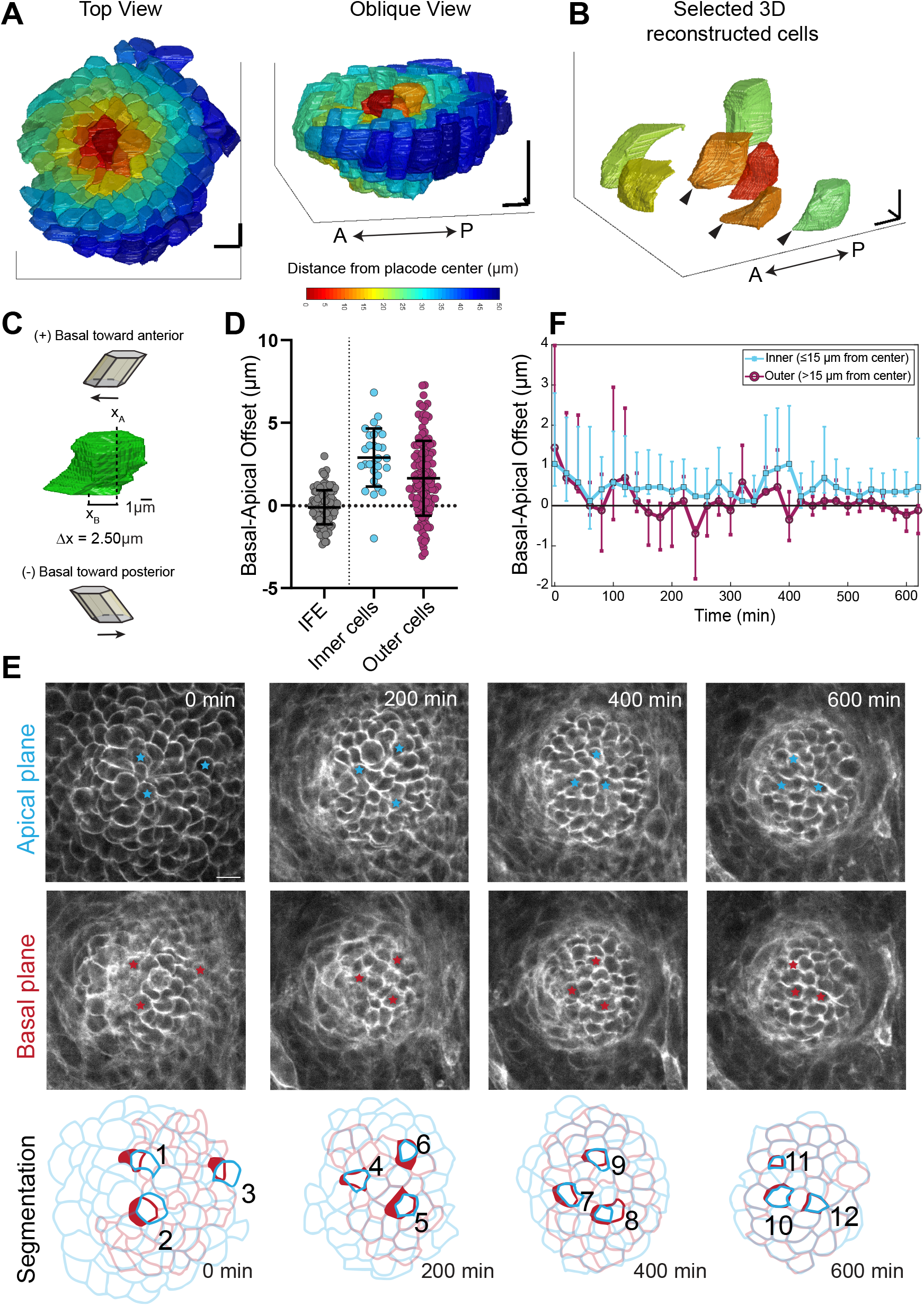
Placode cells form anterior-directed protrusions from their basal surfaces during polarization. A) 3D reconstruction of a polarizing hair placode from *E*15.5 *K14-Cre; mTmG* embryo shown in top and oblique views. Cells are colored based on their distance from center of placode. Scale bars: 10 µm. B) Selected examples of 3D reconstructed inner cells (red and orange, <15 µm from placode center) and outer cells (yellow and green, >15 µm from placode center). Cells are shown at their positions relative to the placode center. Apical is up. Black arrowheads point to anterior-directed protrusions emanating from basal side of inner cells. Scale bars: 5 µm. C) Basal-Apical Offset measurement. 3D reconstruction of a single placode cell (green) showing positions *x*_*A*_ and *x*_*B*_, the x-components of the centroids of the apical and basal surfaces. Basal-Apical Offset is the difference between *x*_*A*_ and *x*_*B*_. Positive offset values indicate the basal surface is more anterior to apical (top schematic). Negative offset indicates the basal surface is more posterior to apical (lower schematic). D) Quantification of Basal-Apical Offsets calculated from 3D reconstructions of cells from the interfollicular epidermis (IFE, grey), inner cells (cyan), or outer cells (magenta) of the polarizing placode. E) Representative time-lapse images of polarizing hair placode from *E*15.5 skin explant expressing membrane-tdTomato. The apical (top panels) and basal (center panels) planes of the placode are shown and were tracked and correlated through time. Blue and red stars mark selected cells highlighted in segmentation masks below. Segmentation masks (bottom panels) for apical (blue) and basal (red) planes are overlayed. Numbered cells highlight different cells at each time point, showing apical and basal segmentations. Anterior to left. Scale bars: 10 µm. F) Quantification of Basal-Apical Offset calculated at each 20-minute timepoint for all placode cells segmented over 10 hours. Cells are classified based on their distance from the placode center, inner ≤15µm (blue) and outer >15 µm (magenta). Each point represents median Basal-Apical Offset per timepoint, and error bars represent first and third quartile.

Since 3D reconstructions from fixed epidermal samples capture just a snapshot of placode cell morphology, we performed live imaging of developing placodes to test whether the basal surfaces of inner cells are consistently anteriorly directed throughout polarization. From time-lapse, confocal z-stacks spanning the full apical-to-basal thickness of the polarizing placode, we extracted the apical- and basal-most planes (Fig. 2E, top and center rows, Movie S5), segmented cell edges, and then paired the apical and basal surfaces of each cell for each timepoint (Fig. 2E, bottom row). At each timepoint during placode polarization, we found that the basal surfaces of many cells were more anteriorly displaced than the apical surface. To quantify this, we calculated the Basal-Apical Offset for all segmented cells at each timepoint, binning cells according to their distance from the placode center (Fig. 2F). Here, any cell within 15 µm of the placode center was classified as an inner cell, and any cell outside of 15 µm from the placode center was classified as an outer cell. This classification is based on previous measurements of the distribution of inner and outer cells within the placode (Leybova et al., 2024). We found that the median Basal-Apical Offset of inner cells was consistently positive (basal anteriorly displaced) over the course of placode polarization. Conversely, the median Offset of outer cells hovered around zero. Thus, anterior displacement of the basal surface in inner cells persists over the course of placode polarization, suggesting that polarized crawling by inner cells may be driving the observed counter-rotational cell flows in the placode. Moreover, these data also suggest that inner cells are likely the main force-generating cells driving polarization, while outer cells likely play a more passive role. This corroborates previous results in mutant embryos showing that loss of inner but not outer fates blocks placode cell rearrangements (Leybova et al, 2024).

### Celsr1 is required for the formation of basal protrusions

We have shown that as placode epithelial cells move, they extend planar polarized, anterior-directed protrusions from their basal surfaces. We next asked whether the core PCP pathway is required for the formation of anterior-directed basal protrusions. To investigate this, we generated 3D reconstructions of hair placodes from embryos lacking Celsr1 (*Celsr1*^*KO/KO*^; *mTmG)*, the upstream-most core PCP protein (Basta et al., 2023; Basta et al., 2025). In the absence of Celsr1 function, PCP establishment is completely abolished in the skin epidermis, and asymmetric localization of Fz6 and Vangl2, as well as hair follicle polarity, are lost (Basta et al., 2023; Devenport and Fuchs, 2008; Ravni et al., 2009). We found that in contrast to cells of wild-type polarizing placodes, cells of *Celsr1*^*KO/KO*^ placodes were relatively columnar in shape and did not form protrusions from their basal surfaces (compare Fig. 2A-B, Movies S2-3 and Fig. 3A-B, Movies S6-7). *Celsr1*^*KO/KO*^ placode cells displayed an inward curvature, but their orientations were radially symmetric around the placode, rather than directed towards the anterior (Fig. 3B). Calculating the Basal-Apical Offset, we found that Offset values were close to zero for both inner and outer cells in a *Celsr1*^*KO/KO*^ placode, showing that neither cell type had a significant bias for anterior directed protrusions. This contrasts with wild-type polarizing placodes, where inner cells displayed a clear bias of the basal surface toward the anterior (Fig. 3C). These data suggest that both the formation and orientation of basal protrusions require Celsr1 and likely PCP function more broadly.

**Figure 3.**
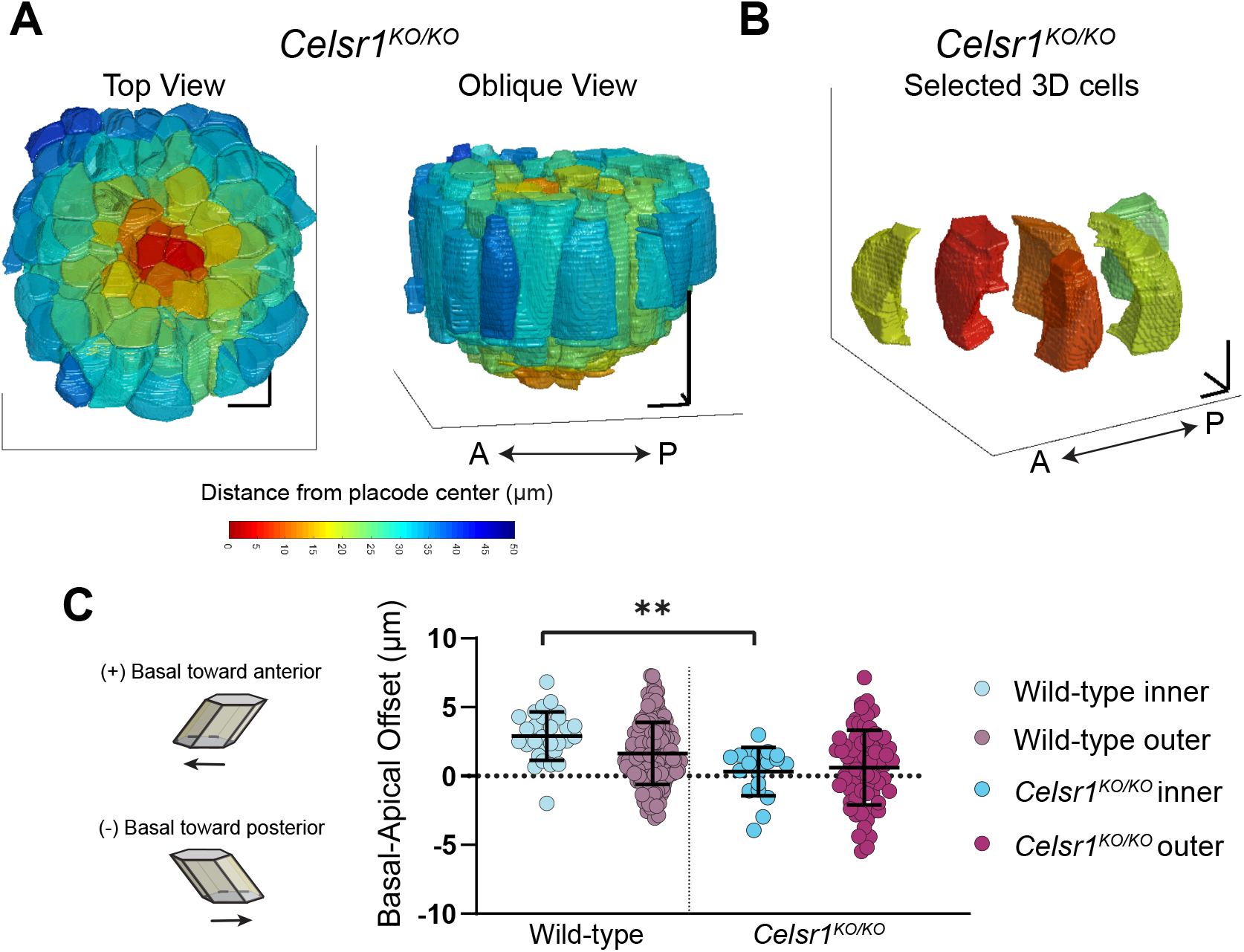
PCP is required for anterior-directed protrusions. A) 3D reconstruction of a polarization-staged hair placode from *E*15.5 *Celsr1*^*KO/KO*^; *mTmG* embryo, shown in top and oblique views. Cells colored based on their distance from the placode center. Scale bars: 10 µm. B) Selected examples of 3D reconstructed inner (red and orange, <15 µm from placode center) and outer cells (yellow and green, >15 µm from placode center). Cells are shown at their positions relative to the placode center. Apical is up. Scale bars: 5 µm. C) Quantification of Basal-Apical Offsets calculated from 3D reconstructions of wild-type polarizing and *Celsr1*^*KO/KO*^ placodes.

### PCP components asymmetrically localize along the basolateral surface of placode epithelial cells

Although PCP proteins are restricted to the apico-lateral region of cell junctions in *Drosophila* (Djiane et al., 2005), in hair placodes, PCP proteins are distributed along the entire lateral surface of adjoining anterior-posterior cell edges (Fig. 4A) (Basta et al., 2021; Devenport and Fuchs, 2008). To investigate whether opposing PCP complexes are indeed partitioned to opposite sides of placode cells at their basolateral edges, we examined Fz6 and Vangl2 localization in chimeric epidermal tissues generated by mixing embryos that co-express Fz6-3xGFP and tdTom-Vangl2 with unlabeled wild-type embryos (*Fz6-3xGFP;tdTom-Vangl2* : *WT* chimera) (Basta et al, 2025). The resulting mosaically-labeled epidermis enabled us to simultaneously visualize opposing PCP components in cells surrounded by unlabeled neighbors. Similar to the surrounding IFE cells, we observed that in placode cells, Fz6-3xGFP was localized posteriorly and tdTom-Vangl2 was localized anteriorly and at both apical and basal surfaces (Fig. 4B). To quantify their asymmetry, fluorescence intensities of Fz6-3xGFP and tdTom-Vangl2 were measured along cell edges, and each cell edge was classified as either anterior or posterior based on its position relative to the cell center. We then calculated the log of the ratio of posterior intensity to anterior intensity for both Fz6-3xGFP and tdTom-Vangl2 and found that Vangl2 was consistently polarized to the anterior edge of cells, while Fz6 was polarized to the posterior edge (Fig. 4C). These data demonstrate that anterior and posterior components of the PCP complex display unipolar, planar polarized localization all along the lateral edges of placode cells. Therefore, PCP proteins are indeed present at the basolateral surface of placode cells, where basal protrusions are observed.

**Figure 4.**
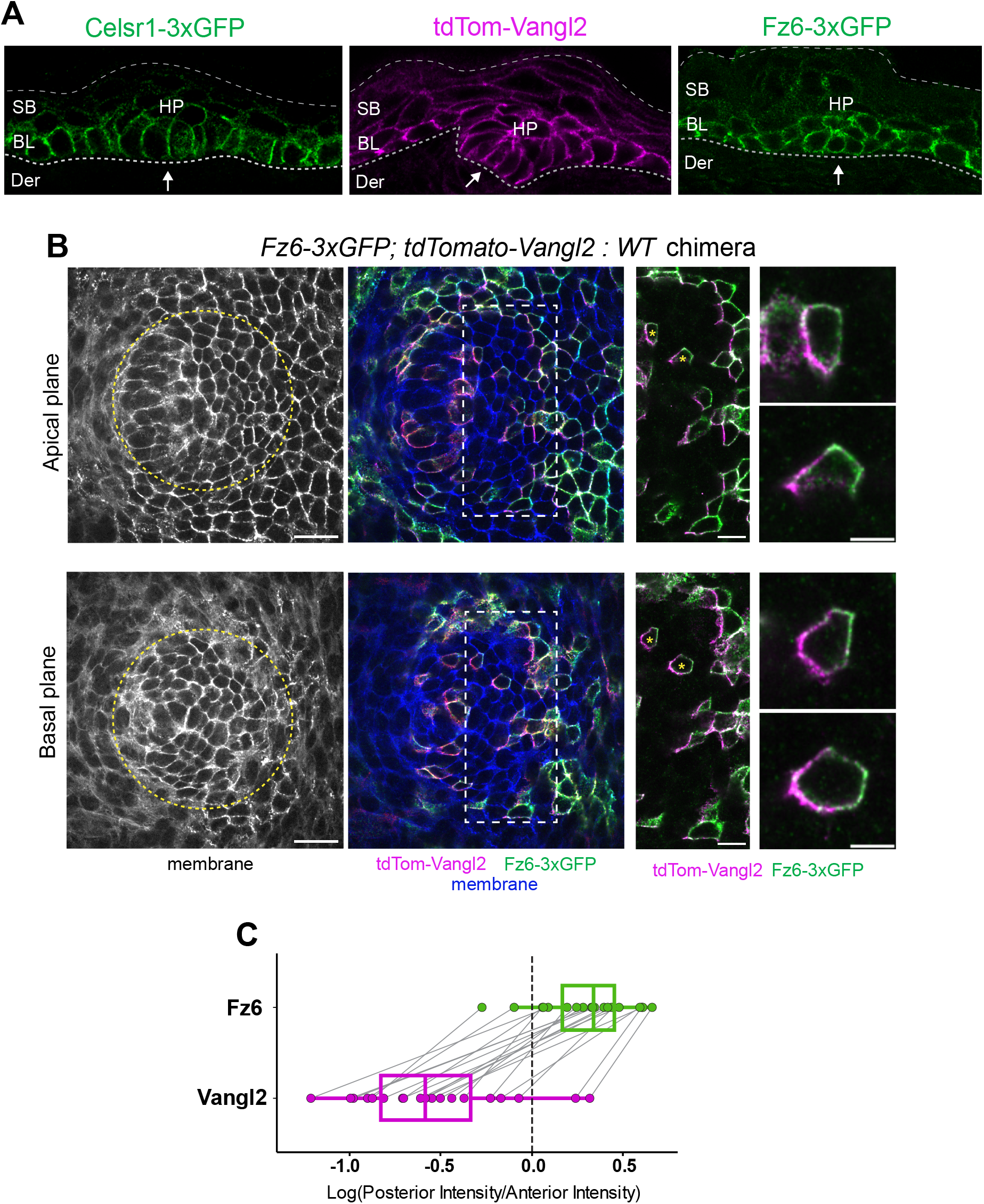
PCP proteins localize along the lateral surfaces of placode cells. A) Representative sagittal views of hair placodes *E*15.5 embryos expressing Celsr1-3xGFP (left, green), tdTom-Vangl2 (middle, magenta), or Fz6-3xGFP (right, green) from their endogenous loci. Lower dotted lines indicate position of the dermal-epidermal boundary. Upper dotted lines depict the upper most suprabasal layer. HP=hair placode, Der=dermis, BL=basal layer, SB=suprabasal layers. Anterior to left. B) Apical plane (top panels) and basal plane (bottom panels) of representative hair placode from *E*15.5 *Fz6-3xGFP; tdTom-Vangl2* : *WT* chimeric skin labeled with phalloidin and Celsr1 to mark membranes. Left panels (grayscale) show membrane label highlighting placode epithelium and dermis. Middle and right panels show mosaic labeling of tdTom-Vangl2 (magenta) and Fz6-3XGFP (green). Yellow asterisks mark selected cells zoomed in on the right. Note the asymmetric localization of PCP proteins at both apical and basal surfaces of the placode. Anterior to left. Scale bars: 20 µm and 5 µm (insets). C) Quantification of anterior/posterior intensity ratio of tdTom-Vangl2 and Fz6-3xGFP at apical surface in chimeric placode cells. Anterior to left.

### Continuum mechanics modeling shows that cell crawling is sufficient to generate apical flows

Our data thus far suggest that cell crawling by inner cells along the extracellular matrix (ECM) may be the driving force behind placode polarization. To test the hypothesis that directed cell crawling at the basal surface is sufficient to generate cell flows apically, we developed a continuum mechanics model of the hair placode, focusing on in-plane deformations that occur during polarization (schematic in Fig. 5A). The mouse epidermis is a multi-layered epithelium, where the basal-most layer consists of the IFE and placode cells; apical to those are suprabasal cells. Basal to the epidermis is the dermis, separated by a basement membrane. The basal layer of the epidermis and the surrounding tissues are modeled as a system consisting of a flat viscoelastic slab situated between two fluid layers and solid hard walls (Fig. 5B) (Du and Shelley, 2023). The viscoelastic slab (region *S*) represents a single hair placode and the surrounding IFE cells. As we are interested in the early stages of placode polarization, we assume that the placode remains flat and does not buckle out of plane. The placode and IFE cells are modeled as an Oldroyd-B viscoelastic fluid, a continuum model derived from the coarse-graining of spring-like polymers embedded in a viscous fluid (see Supplemental Methods). To nondimensionalize our model equations, we rescale time by the relaxation time of the Oldroyd-B polymers *τ* = *τ*_*p*_, length *ℓ*_0_ by the radius of a typical placode cell (∼4 µm), and force by a force scale *F*_0_ that is proportional to the viscosity of the tissue *η*; that is, 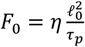. Following the nondimensionalization procedure, our parameters become dimensionless.

**Figure 5.**
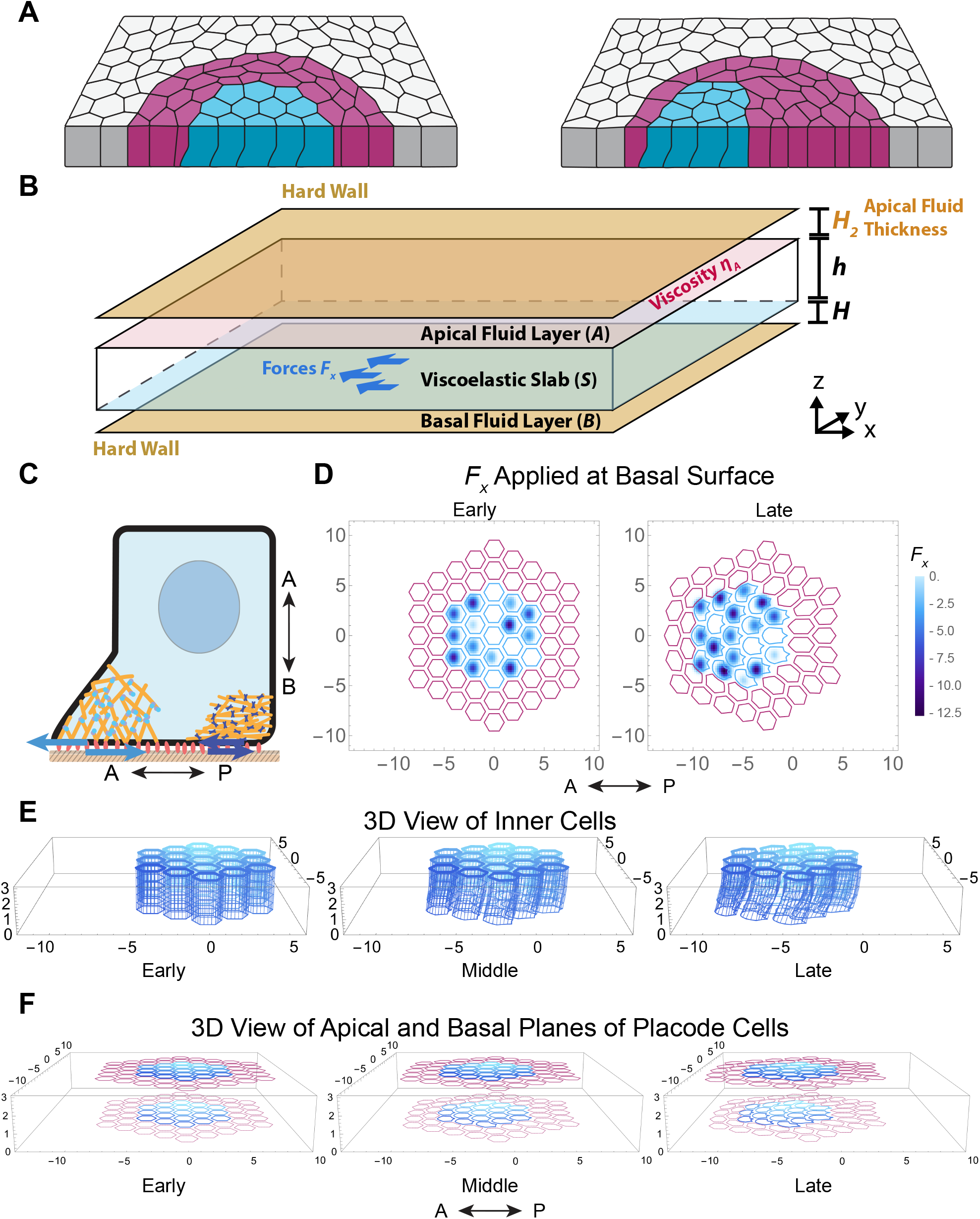
Model of hair placode polarization shows that cell crawling forces applied at basal surface leads to experimentally observed slanted cell shapes. A) Schematic of hair placode focusing on in-plane deformations at the onset of polarization (left) and during polarization (right). The placode is surrounded by IFE cells (gray). Inner cells (cyan) crawl towards the anterior, displacing outer cells (magenta). B) Schematic for continuum model of the hair placode. Viscoelastic Slab (region *S*) represents placode cells and surrounding IFE. Apical Fluid Layer (region *A*) represents apical adhesion interface between placode cells/IFE and overlying suprabasal cells. The apical stationary hard wall with a no-slip boundary condition represents stationary suprabasal cells. Strength of adhesion is modeled by viscosity *η*_*A*_. Basal Fluid Layer (region *B*) represents the basal adhesion interface between placode cells and the underlying basement membrane and dermis. The basal stationary hard wall with a no-slip boundary condition represents the stationary basement membrane. Crawling forces *F*_*x*_ by inner cells are applied at the interface between region B and region S (blue arrows). *H*_2_, *h*, and *H* refer to the thickness of regions *A, S*, and *B*, respectively. C) Schematic of forces to model cell crawling. The cell is attached to the underlying basement membrane by proteins such as integrins (red ovals). Actin cytoskeleton (yellow) attaches to integrins through a variety of intermediate proteins (not shown). At the anterior of the cell, branched actin forms through Rac1-mediated polymerization, involving proteins complexes like WAVE and Arp2/3 (blue and purple dots). This generates anterior directed protrusions, as a result of a force dipole (light blue arrows). At the posterior of the cell, myosin (purple), linking filamentous actomyosin, generates a contractile rear force, resulting in a force dipole at the rear of the cell (dark blue arrows). As we treat the basement membrane to be anchored, we assume the rear-ward halves of each force dipole is absorbed by the basement membrane and is therefore not modeled. D) Plot depicting how forces are simulated at the basal surface of region *S*, shown in planar view. Each inner cell independently applies crawling forces by alternating anterior and posterior patterns of force. Example fields *F*_*x*_, the *x*-component of the force, at early (left) and later (right) timepoints of the simulation are shown. Values of *F*_*x*_ are negative to model force towards the anterior. See also Movie S8 (left). See Supplemental Methods for full implementation details. E) 3D view of tracked points in simulation of inner cells throughout the thickness of region *S* over the course of simulation time (left to right). See also Movie S9 (center). F) 3D view of surface points in simulation tracked for inner and outer cells over the course of simulation time. See also Movie S8 (right). Anterior is to the left, apical is up for all simulations.

We model the sheetlike epidermis as infinitely periodic in the *x* and *y* directions while having a finite thickness in the *z* direction. Our description does not contain individual cells, as we are focused on deformations at the multicellular scale. Since the tissue is modeled as a continuum, we track points embedded in the fluid that move via the velocity field; these points are initially arranged in a pattern of vertical hexagonal prisms, each of which represents the initial shape of a cell. As the simulation progresses and these tracked points move, their new positions represent the deformed shape of a cell. Hexagonal prisms initiated near the center of the simulation domain represent the inner cells while the surrounding hexagonal prisms represent the outer cells. The region surrounding the outer cells represents the IFE, where we do not track points.

The apical fluid layer (region *A*) represents the apical adhesion interface between the placode cells and the overlying suprabasal cells. The apical hard wall corresponds the bulk of the suprabasal layer that remains static during placode polarization. The basal fluid layer (region *B*) represents the adhesion interface between the basal surface of the placode and IFE cells and the underlying basement membrane. The basal hard wall represents the basement membrane itself, which resists motion. Note that both hard walls represent the point where no movement occurs during the early stages of placode polarization, as we have not observed significant deformations within either the suprabasal cells or the basement membrane. We estimated the thickness of the fluid layers from measurements by confocal and electron microscopy: the thickness of region *A* (*H*_2_) is ∼0.2-2 µm and the thickness of region *B* (*H*) is ∼0.5 µm (Jones et al., 2023).

Since we hypothesized that inner cell crawling provides the driving force for placode polarization, in the model, we specify active forces from only the inner cells. Outer and IFE cells remain passive. Cell crawling in our model is prescribed based on findings in lamella-based migration, where cells are front-rear polarized, and crawling occurs as cycles of protrusions at the front followed by contractions at the rear. We implement crawling forces as monopoles, since protrusive and contractile machinery are anchored to the basement membrane, assumed to be rigid. Hence, the force that the cell applies to the basement membrane is not modeled. Forward force at the front occurs through actin polymerization driving protrusions, while forward force at the rear occurs through myosin motors contracting actin filaments (Fig. 5C). We therefore model crawling as phases of anterior-directed forces in the anterior half of a cell, followed by anterior-directed forces in the posterior half of a cell (Fig. 5D and Movie S8). Details on force implementation are in the Supplemental Methods.

We found that within a wide range of parameters, simulations broadly resembled the early stages of placode polarization. Specifically, while cell shapes in the model were initialized to be columnar, as the simulation progressed, the basal surface of cells became more anteriorly displaced compared to their apical surface, similar to confocal data (compare Fig. 5E and Fig. 2B). Although, we did not observe T1 transitions, this is not unexpected, as modeling neighbor exchanges may require specifying local junctional forces or forces due to cell cortex rigidity (Du and Shelley, 2023). Our observations show that a 3D continuum description can serve as a useful way to model the transmission of forces from one surface of the tissue to another.

We then explored model parameters that give rise to different cell shapes and patterns of motion (Movie S9). Specifically, we modulated the magnitude of the active force for cell crawling (*F*_*x*_) and the viscosity of the apical fluid layer (*η*_*A*_). The force *F*_*x*_ is of interest since it varies greatly between different migrating cell types and is unknown for placode cells. Meanwhile, *η*_*A*_ models how tightly placode cells adhere to the overlying suprabasal layer; as the two layers express different cadherins, the strength of the apical adhesion interface (*η*_*A*_) might be regulated (Jamora et al., 2003; Leybova et al., 2024; Müller-Röver et al., 1999). Finally, we varied *H*_2_ to explore how the thickness of the apical fluid layer can contribute to cell flows. We explored *H*_2_ because it was recently shown that during polarization, the apical surfaces of outer cells bend inward to form a canopy above inner cells (Leybova et al., 2024), potentially altering the thickness of the adhesive interface.

We performed simulations varying apical adhesion interface parameters *η*_*A*_ and *H*_2_ (fixing *F*_*x*_). Simulations gave rise to a variety of patterns of motion. To visualize this motion, we obtained velocity fields at the apical- and basal-most surfaces of the model placode. We subtracted out velocities such that the average motion at the northernmost and southernmost edges of the IFE was zero, allowing us to examine relative motion within the placode, rather than the overall anterior-ward movement. Figure 6A and Movie S10 show velocity vector fields from two dimensionless parameter sets with the same *F*_*x*_, one with low *η*_*A*_ = 0.05 and moderate *H*_2_ = 0.5 (Fig. 6A left column) and the other with high *η*_*A*_ = 2.0 and low *H*_2_ = 0.05 (Fig. 6A right column). Both simulations resulted in similar velocity patterns at the basal surface, showing predominantly anterior-ward motion (Fig 6A, bottom row). In contrast, velocities at the apical surface for the first simulation (low *η*_*A*_, moderate *H*_2_) displayed counter-rotational, vortical flows (Fig. 6A, middle and top left), whereas apical velocities for the second simulation (high *η*_*A*_, low *H*_2_) displayed nearly uniform anterior-ward flow (Fig. 6A, middle and top right). To visualize the vorticity of these two apical flows, we calculated the curl of the apical velocity field (vorticity). Here, large absolute values correspond to more vortical flows, negative values correspond to clockwise flow, and positive values correspond to counterclockwise flow. In the first simulation with strong vortical flow, we observed two clear zones of high vorticity in opposing directions above and below the midline of the placode (Fig. 6B and Movie S11). This contrasts with the second simulation with nearly uniform flow, where we observed that vorticity was greatly decreased (Fig. 6B and Movie S11). Our continuum modeling shows that altering parameters associated with the geometry and material properties of the apical adhesion interface can affect the formation of vortical velocities. Moreover, we show that basal crawling forces are sufficient to generate apical flows in a counter-rotational pattern.

**Figure 6.**
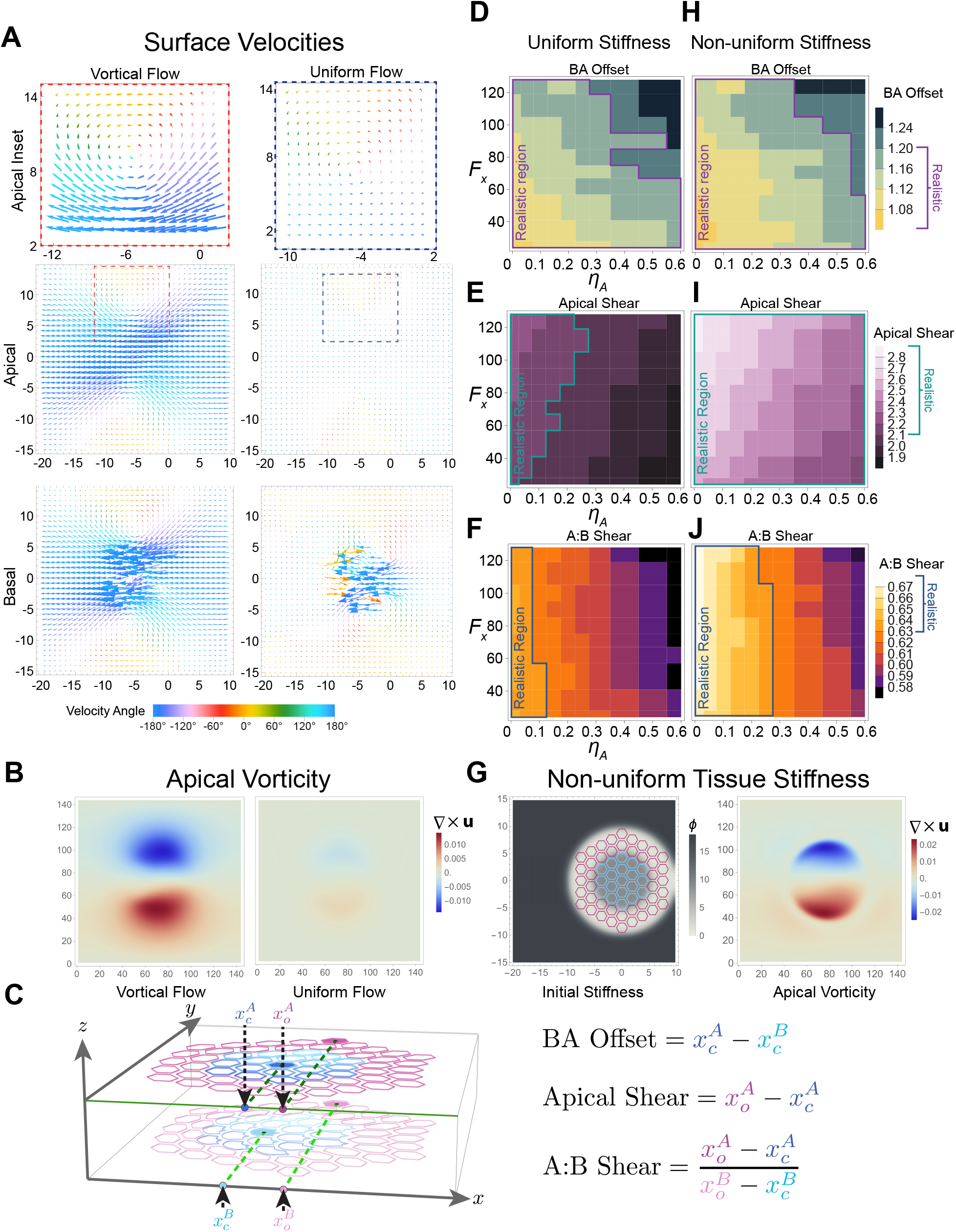
Lower apical adhesion and non-uniform tissue stiffness promote formation of apical counter-rotational flow in model. A) Apical velocity fields of the apical (middle panel) and basal (bottom panel) surfaces of the model slab comparing two parameters sets showing vortical (left) and uniform (right) flow at the apical surfaces. Zoomed-in insets of the northern region of apical velocity field for both parameter sets are shown (top panel). Vector length is proportional to velocity magnitude and colored based on angle from -180° to 180°. For both simulations, *F*_*x*_ = 64. Left: *H*_2_ = 0.5, *η*_*A*_ = 0.05. Right: *H*_2_ = 0.05, *η*_*A*_ = 2.0. B) Vorticity fields computed as the curl of apical surface velocity at late simulation times (once central cell reaches *x* = −6) for simulations shown in (A). Negative values indicate clockwise vortical flow (blue), and positive values indicate counterclockwise vortical flow (red). C) Schematic depicting metric calculation from simulation results. Metrics are calculated from four distances measured in each simulation. The values 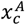 and 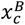 represent the distance traveled by the centroid of the central cell (shaded blue cell) along the x-axis at the apical and basal surfaces, respectively. Similarly, 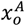 and 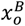 represent the distance traveled by the centroid of the northern-most outer cell (shaded pink cell) along the x-axis at the apical and basal surfaces, respectively. D-F, H-J) Phase diagrams of Basal-Apical Offset (D, H), Apical Shear (E, I), and A:B Shear (F, J) measured from simulations with *η*_*A*_ = 0.001 to 0.6 (x-axis) and *F*_*x*_ = 24 to 128 (y-axis). Diagrams correspond to simulations with uniform tissue stiffness *ϕ* = 9 (D-F), and non-uniform tissue stiffness *ϕ*_*IFE*_ = 18, *ϕ*_*Outer*_ = 1, *ϕ*_*Inner*_ = 9 (H-J). *Realistic* metric regions defined as simulations which resulted in metric values in the following ranges: BA Offset 0.25-1.20, Apical Shear greater than or equal to 2.1, and A:B Shear 0.63-1.0. G) Non-uniform tissue stiffness implemented by non-uniform density of Oldroyd-B particles *ϕ*. Scalar field *ϕ* specified based on initial positions of inner, outer, and IFE cells (left). Resulting apical vorticity field from surface velocity of non-uniform stiffness simulation at late simulation time (right).

To compare many different simulations with varying parameter values, we developed three metrics to quantify characteristics of cell shape and flow pattern (Fig. 6C). These metrics were calculated using the displacement along the x-axis (*x*) of the central inner cell and the northernmost outer cell (subscript *c* and *o*, respectively). These *x*-displacements are measured at both the apical and basal surfaces (superscript *A* and *B*, respectively). We define the metrics Basal-Apical Offset (BA Offset), Apical Shear, and Apical:Basal Shear (A:B Shear):

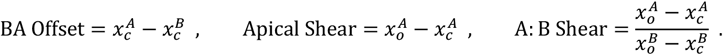

Here, BA Offset is similar to the BA Offset metric analyzed for experimental data in Fig. 2C-F, 3C. Correspondingly, a more positive BA Offset results from a greater anterior displacement of the basal surface. Apical shear describes the difference in the distance traveled by the center cell compared to the northernmost cell at the apical surface. A greater Apical Shear corresponds to a stronger vortical pattern of flow at the apical surface. Finally, the A:B Shear is a metric for the similarity between the apical and basal vorticity, as it computes the ratio between the Shear at the apical side and the Shear of the basal side. An A:B Shear close to 1 results from little difference between apical and basal flows. Typically in simulations, the A:B Shear is less than 1, as counter-rotational flow is typically weaker at the apical surface. All metrics are computed when the central inner cell reaches position *x* = −6. This corresponds to the center cell of the placode having traveled ∼24 µm along the AP axis, consistent with the average displacement observed over ∼10 hours during an average time-lapse of hair placode polarization (Fig. 2E).

We ran simulations varying *F*_*x*_ = 24 to 128 and *η*_*A*_ = 0.001 to 0.6. We used *η*_*B*_ = 0.2 for the viscosity of the basal fluid layer, as this layer is expected to be less viscous than the tissue (*η* = 1 from nondimensionalization). We fix *H*_2_ = 0.5, as this moderate value resulted in apical vortical flows at the apical surface (Fig 6A). Based on measurements from experiments, metric values should fall in the ranges of: BA Offset between ∼0.25-1.20, Apical Shear greater than 2.1, and A:B Shear around 0.7-1.0. We defined *realistic parameters* as those that generated simulations with metrics that fell approximately within these *realistic metric* ranges. In Fig. 6D-F, we find that many simulations with realistic metric values are in the parameter space of *η*_*A*_ < *η*_*B*_ = 0.2, and as *η*_*A*_ increases past *η*_*B*_ = 0.2, we see significant changes in all three metrics into regimes outside of realistic values. Interestingly, this result that realistic parameters mostly satisfy *η*_*A*_ < *η*_*B*_ is consistent with experimentally observed relative protein adhesion strengths, as measurements show that cell-basement membrane interactions (basal) are stronger than cell-cell interactions (apical) (Arimori et al., 2021; Buck and Horwitz, 1987; Vendome et al., 2014). Examining the relationship between *F*_*x*_ and each metric, we find that BA Offset strongly depends on *F*_*x*_, Apical Shear only weakly depends on *F*_*x*_, and A:B Shear Ratio is nearly independent of *F*_*x*_. From the BA Offset plot, realistic parameter values can be found for all ranges of *F*_*x*_; however, the range of *η*_*A*_ resulting in realistic metric values is broader at lower *F*_*x*_. If the viscoelastic relaxation time *τ*_*p*_ were known for the hair placode, we could further constrain *F*_*x*_ based on time needed for the tissue to deform to the expected shape. These results show that we are able to simulate the slanted cell shapes and counter-rotational flow consistent with experimental metrics for a sizable range of parameter values.

We further investigated varying the material properties of the placode and IFE. Previous studies have found that spatial differences in tissue stiffness play a role in morphogenesis (Parada et al., 2022; Peak et al., 2022; Petridou et al., 2019; Shyer et al., 2015; Yang et al., 2023). Hence, to test the role of differences in stiffness across the tissue, we performed simulations where the density of Oldroyd-B polymers *ϕ* was spatially non-uniform (Fig. 6G left; see Supplemental Methods for implementation). We assigned the IFE to have high stiffness of *ϕ* = 18 in dimensionless units, inner cells to have moderate stiffness of *ϕ* = 9, and outer cells to have low stiffness of *ϕ* = 1. This contrasted with previous simulations with uniform stiffness where *ϕ* = 9 everywhere. In the non-uniform stiffness simulations, we found two zones of strong vorticity with larger magnitudes compared to those of uniform stiffness simulations (compare Fig. 6B left and 6G right). Additionally, we observed that high vorticity was restricted to a circular region by the stiffer surrounding IFE (Fig. 6G right). Exploring the parameter space of *F*_*x*_ and *η*_*A*_ to compare uniform and non-uniform stiffness simulations (Fig. 6D-F and 6H-J), we see a significant increase in the range of realistic parameters when stiffness is non-uniform. Based on these results, we hypothesize that confinement by a highly stiff IFE together with softer outer cells allows the inner cells to displace the outer cells more effectively, creating a stronger counter-rotational flow. Our model shows that non-uniform tissue properties could increase the vortical velocity at the apical surface due to basal forces and may be one way for the placode to increase robustness in polarization.

### Rac1 is required for hair placode polarization

Given that basal crawling forces are sufficient to generate the movements associated with placode polarization in our computational model, we next asked whether cell crawling is necessary for polarization *in vivo*. Cells migrating along an ECM typically display front-rear polarity with a protrusive front and a contractile rear (Campellone and Welch, 2010; Insall and Machesky, 2009; Ridley, 2011). At the front, lamellipodia extensions are initiated by Rac1-dependent branched actin, while rear retractions form through RhoA-dependent actomyosin contractility (Vicente-Manzanares et al., 2009). In previous work, we showed that placode polarization requires contractile actomyosin (Cetera et al., 2018), but it remains unknown whether leading edge regulators such as Rac are also required.

To determine whether Rac activity is required for hair placode morphogenesis, we cultured *E*15.5 *Fz6-3xGFP; mTomato* skin explants with the Rac1 inhibitor NSC 23766. Following 22 hours culture in varying concentrations of NSC 23766 (10, 25, or 50 µM), explants were fixed and labeled for Sox9, a marker of outer cells, which are normally displaced posteriorly during placode polarization (Fig. 7A). Fz6-3xGFP localization remained planar polarized in explants cultured with NSC 23766 (Fig. S3). Using the posterior:anterior ratio of Sox9 fluorescence intensity as a measure of placode polarity, we observed a dose-dependent inhibition of placode polarization upon treatment with NSC 23766 (Fig 7A-C). In the presence of 25 or 50 µM, but not 10 µM NSC 23766, placodes that had been specified but not yet polarized at the start of inhibitor treatment completely failed to polarize, such that Sox9-expressing cells remained localized in rings around inner cells (Fig. 7A,C).

**Figure 7.**
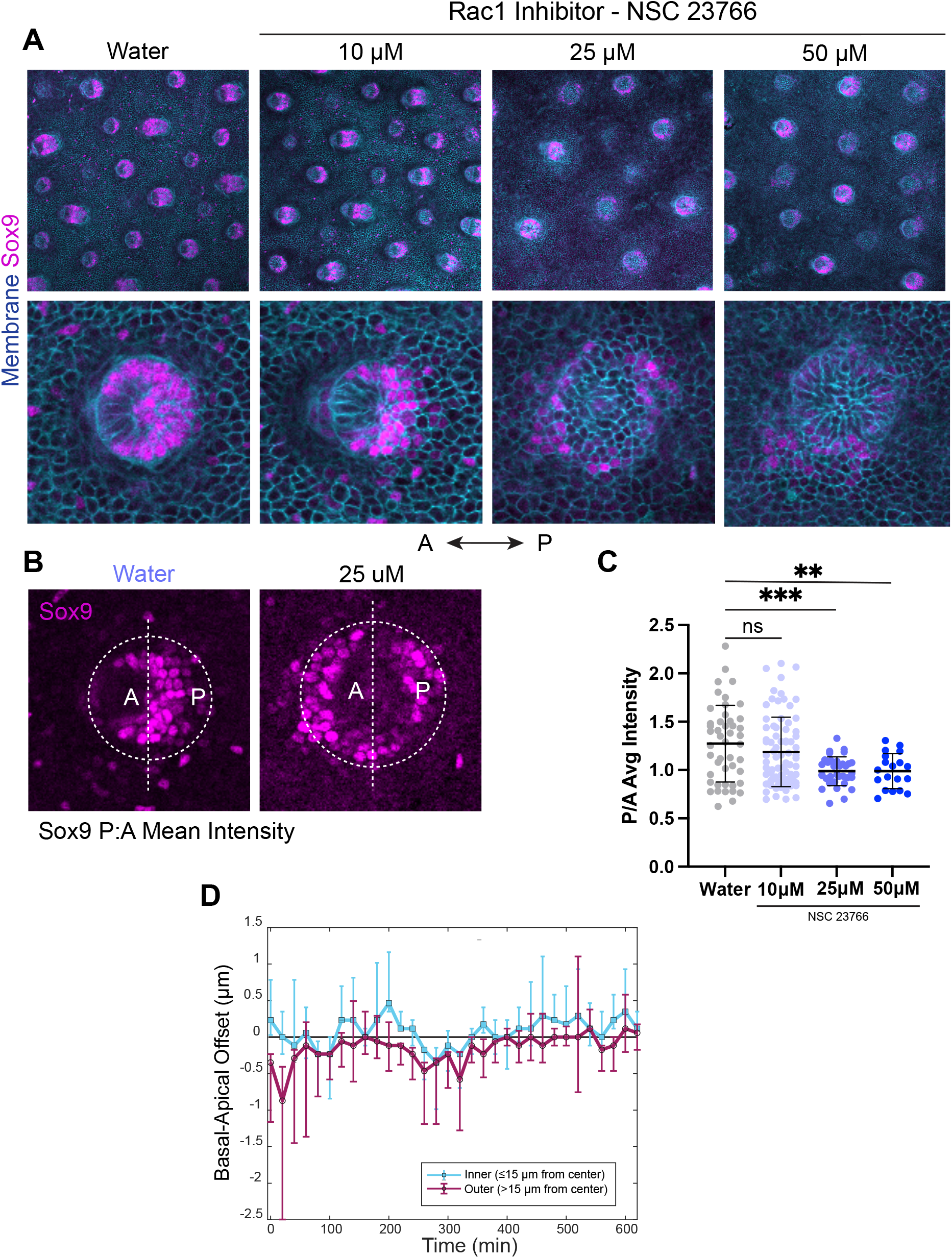
Inhibition of Rac activity prevents anterior-directed migration and polarization of hair placodes. A) Representative planar views of *E*15.5 mTmG skin explants cultured for 22 hours with increasing concentrations of Rac inhibitor NSC 23766 or water as a control. mT (cyan) and Sox9 (magenta), which labels outer cells of pre-polarized and posterior cells of polarized placodes, are shown. Top panels display zoomed out view containing several hair placodes at different stages of development (first, second and third waves of induction; Scale bars: 100 µm). Bottom panels show high magnification views of individual second-wave placodes that would have undergone polarization during the culture period (Scale bars: 10 µm). Note that third wave placodes fail to form in explants treated with 25 µM or 50 µM NSC 23766. B) Method for measuring hair placode polarization in (C) using the ratio of mean Sox9 intensity on the posterior versus anterior side of the placode. C) Quantification of hair follicle polarization (posterior:anterior average Sox9 intensity) in control and NSC 23766-treated explants. All three waves of hair placode induction are included for each group. D) Quantification of Basal-Apical Offset of placode cells treated with 50 µM NCS 23766 from time-lapse imaging calculated at each 20-minute timepoint for all placode cells segmented over 10 hours. Cells are classified based on their distance from the placode center, inner ≤15µm (blue) and outer >15 µm (magenta). Each point represents median Basal-Apical Offset per timepoint, and error bars represent first and third quartile.

To investigate whether the loss of placode polarization upon Rac1 inhibition was due to a defect in anterior-directed cell crawling, we performed live imaging of NSC 23766-treated *Fz6-3xGFP; mTomato* skin explants and calculated the Basal-Apical Offset of placode cells over a 10-hour time period. In contrast to untreated controls, in which inner cells displayed median Basal-Apical Offset values greater than zero at all timepoints, and were thus anterior-directed, the median inner cell Offset values of Rac1-inhibited placodes oscillated around zero and were therefore undirected (compare Fig. 2F and 7D). These results indicate that Rac1 is required for anterior displacement of the basal surface of inner cells. Moreover, placode polarization requires Rac1-dependent crawling movements, consistent with the idea that crawling along the ECM could serve as the main force driving placode polarization.

## Discussion

The mechanisms driving cell intercalation have been broadly categorized as contraction-based or crawling-based (Shindo, 2018). In the contraction mode, cell intercalation occurs through polarized myosin contractility at apical junctions, leading to orientated neighbor exchanges across a tissue (Bertet et al., 2004; Nishimura et al., 2012; Shindo and Wallingford, 2014; Zallen and Wieschaus, 2004). Conversely, in crawling mode, intercalation occurs through the formation of migratory protrusions and adhesion with neighboring cells and the ECM (Davidson et al., 2006; Keller, 2002; Keller and Tibbetts, 1989; Keller et al., 1992). As epithelial cells adhere tightly to one another whereas mesenchymal tissues consist of more loosely connected migratory cells, these distinct modes of intercalation had been the prevailing models for epithelial and mesenchymal cell rearrangements, respectively. However, it is now clear that epithelia and mesenchymal cells can employ both modes of intercalation. In *X. laevis* gastrulation, contraction-based junction remodeling integrates with cell protrusions to drive neighbor exchanges in the mesoderm, a mesenchymal tissue (Huebner and Wallingford, 2018; Shindo and Wallingford, 2014; Weng et al., 2022). Conversely, basal crawling-like behaviors are observed in epithelial intercalation including *C. elegans* dorsal hypodermis convergent extension (Williams-Masson et al., 1998), *Drosophila* germband extension (Sun et al., 2017), ascidian notochord formation (Munro and Odell, 2002), and mouse neural tube closure (Williams et al., 2014). It is interesting to note that early studies investigating epithelial neighbor exchanges suggested a ‘cortical tractoring’ mechanism during neural tube closure, where the basal and lateral, not apical, surfaces were the sites of force generation (Cheng et al., 1987; Jacobson and Gordon, 1976; Jacobson et al., 1986; Moury and Jacobson, 1989). Our study, showing basal crawling of placode epithelial cells undergoing neighbor exchange, lends further support to the idea that the basal surface can be the site of force generation in epithelial morphogenesis.

Critical to this model is that basal forces ultimately result in neighbor exchanges observed at the apical surface of the cell. As demonstrated by our continuum model of the hair placode, forces applied at the basal surface are sufficient to generate flows at the apical surface, as well as medial-lateral vortices, akin to the experimentally observed counter-flow by outer cells. Our model further showed how the correspondence of apical and basal flows is affected by material properties of the tissue. Changes in the apical adhesive interface greatly affected apical flow, in which low viscosity allowed strong correspondence with the basal flow and the formation of vortices, and a viscous apical interface did not. Previous studies have shown that this apical adhesion interface is indeed regulated during placode polarization, both at the protein and tissue level. Placode cells and their overlying suprabasal cells express different classical cadherins, with P-cadherin elevated in the placode and E-cadherin expressed in suprabasal cells (Müller-Röver et al., 1999; Tinkle et al., 2004). Additionally, outer cells form a canopy above inner cells, reducing the contact between inner cells and the overlying suprabasal cells (Leybova et al., 2024). In line with this past work, our continuum modeling suggests that reducing inner cell adhesion to surrounding cells may be required for tissue flows during placode polarization.

Spatial differences in tissue stiffness are known to contribute to morphogenesis (Parada et al., 2022; Peak et al., 2022; Petridou et al., 2019; Shyer et al., 2015; Yang et al., 2023), and our model illustrates the importance of non-uniform material properties in placode polarization. In otherwise identical simulations, regimes where tissue stiffness was non-uniform between inner, outer, and IFE cells had a larger space of realistic parameters compared to uniform stiffness. Thus, one prediction from the model is that material properties may be regulated in the placode to increase robustness of polarization, in which inner, outer, and IFE cells develop different stiffnesses as part of their distinct cell fate identities. Interestingly, opposing, radial gradients of E- and P-cadherin are expressed across the placode epithelium, and disruption of the differential adhesion gradient results in delayed cell flows (Jamora et al., 2003; Leybova et al., 2024; Müller-Röver et al., 1999). In addition, outer cells are more proliferative than inner cells (Ahtiainen et al., 2014; Ouspenskaia et al., 2016), a property associated with increased tissue fluidity both theoretically and experimentally (Firmino et al., 2016; Ranft et al., 2010). Whether these differences in adhesion and proliferation result in changes to tissue stiffness in the placode requires further investigation. Additionally, we observed enhanced counter-rotational flows in simulations where stiffness was increased in the IFE, confining flows in a circular pattern. Recently, it was shown that confinement from the surrounding epidermal tissues and a dermal ring of fibroblasts is required for hair placode downgrowth (Villeneuve et al., 2024). Our model suggests that this confinement may also enhance the counter-rotational flow during placode polarization.

For basal crawling forces to result in stereotyped, directional tissue movement, they must be coupled to cell polarity. We propose that the core PCP system establishes a front-rear polarity in each migratory cell of the placode, aligning the direction of migration with the anterior-posterior axis. In models of cell crawling in both single and collectively migrating cells, Rac1 acts at the front of the cell to generate branched actin-based lamellipodia, while in the rear, RhoA promotes actomyosin-based contractions (Artemenko et al., 2014; Devreotes and Horwitz, 2015; Lauffenburger and Horwitz, 1996; Neumann et al., 2018; Shi et al., 2013; Sun et al., 2017; Williams et al., 2022; Zhu and Nelson, 2013). Here, we showed that Rac1 is required for hair placode polarization, and in previous work, we found that RhoA and Myosin activities were required (Cetera et al., 2018). Together with other studies on actin regulators, these findings indicate that regulation of both branched actin as well as contractile actomyosin are required for hair placode polarization (Cohen et al., 2019; Laurin et al., 2019). Notably, the Fat2-Lar planar polarity system in flies has been shown direct front-rear migration and basal crawling during *Drosophila* follicle cell rotation of the egg chamber (Barlan et al., 2017; Cetera et al., 2014; Williams et al., 2022). We propose the core PCP system could orient migration similarly, through Vangl2-dependent protrusions and Fz6-coupled contractions, providing a new avenue to study mechanisms of planar-polarized collective cell movements.

## Acknowledgements

We would like to thank Drs. Stas Shvartsman, Ricardo Mallarino, Michael J. Shelley, and members of the Devenport Lab for helpful discussion and insights. We would like to thank Gary Laevsky and Sha Wang of the Confocal Imaging Core Facility at Princeton University, a Nikon Center of Excellence, for assistance with imaging. We would also like to thank Katie Little for assistance with mouse husbandry and breeding. The Flatiron Institute is a division of the Simons Foundation. R.S. thanks the CCB at the Flatiron Institute for hospitality while a portion of this research was carried out. The computations reported in this paper were in part performed using resources made available by the Flatiron Institute. We thank Drs. Ann E. Sutherland and late Robert S. AdeIstein for the Myosin IIB-GFP mice. This work was supported by the NSF-GFRP awarded to R.S. Work in the Devenport lab is supported by NIH-NIAMS R01AR066070, NIH-NIAMS R01AR068320 and NIH-NICHD R01HD105009. L.L. was supported by NIH-NIGMS under award number T32GM007388 and by NIH-NIAMS under award number F31AR074246. A.S. was supported by NIH-NIGMS under award number T32GM148739.

## Author Contributions

R.S. was involved in conceptualization, formal analysis, investigation, methodology, software, writing and editing, and visualization. X.D. was involved in conceptualization, investigation, methodology, resources, software, supervision, writing and editing, and visualization. L.L. was involved in investigation and formal analysis of wild-type 3D reconstructions. A.S. was involved in formal analysis of Rac1 timelapse analysis. A.B. contributed to methodology and software for generation of 3D volumetric reconstructions. D.D. contributed to the conceptualization, methodology, writing and editing, visualization, and supervision of the study and was responsible for funding acquisition.

## Methods

### Mouse lines and breeding

All procedures involving animals were approved by Princeton University’s Institutional Animal Care and Use Committee (IACUC). Mice were housed in an AAALAC-accredited facility in accordance with the Guide for the Care and Use of Laboratory Animals. This study was compliant with all relevant ethical regulations regarding animal research. E15.5 embryos from mixed backgrounds were used. Both sexes were used as sex was not determined in embryos. Experiments were performed using several mouse lines: Myh10^tm8Rsad^ (MGI:5518638, *Myosin IIB*^*GFP/GFP*^), Celsr1^*em1Ddev*^ (MGI:7434998, *Celsr1*^*KO/KO*^), Fzd6^*em1Ddev*^ (MGI:7382794, *Fz6*^*3xGFP/3xGFP*^), Vangl2^*em1Ddev*^ (MGI:7382795, *Vangl2*^*tdTom/tdTom*^), Tg(KRT14-cre)1Efu (MGI:1926500, *K14-Cre/+*), and Gt(ROSA)26Sor^tm4(ACTB-tdTomato,-EGFP)Luo^ (MGI:3716464, *Rosa26*^*mTmG/mTmG*^). Embryo genotypes used for each experiment are listed by Figure in the table below.

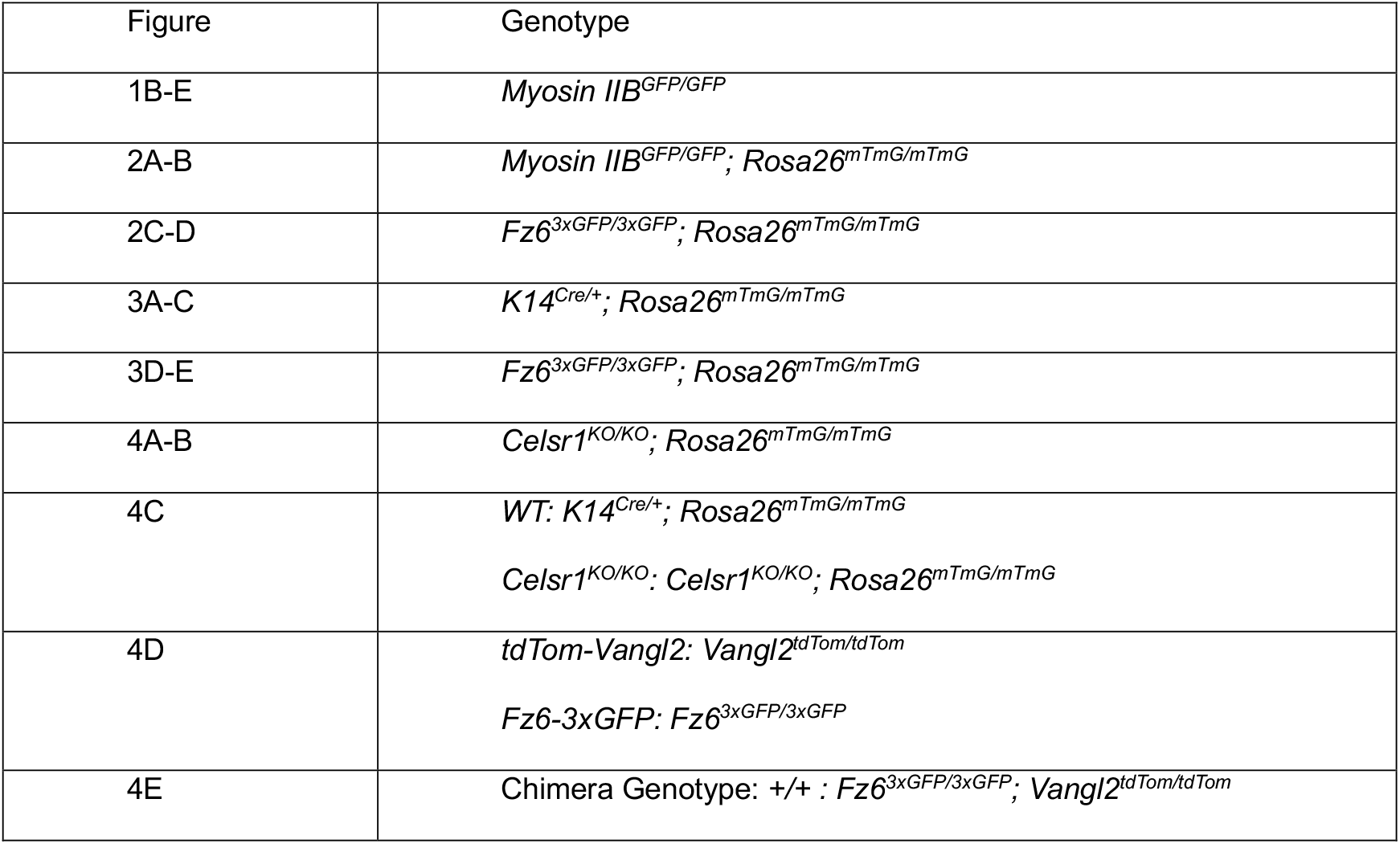
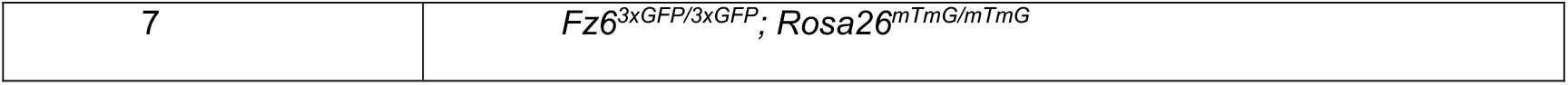

### Whole-mount immunostaining

E15.5 embryos were dissected in PBS with Ca2+ and Mg2+ (PBS++) and fixed in 4% paraformaldehyde/PBS++ for 1 hour and then washed in PBS at room temperature. Back skins were then dissected and blocked overnight at 4°C. Blocking solution consisted of 2% normal goat serum, 2% normal goat serum, 1% bovine albumin, 1% fish gelatin in PBT2 (0.02% Triton in PBS). Primary antibodies were added to blocking solution and incubated overnight at 4°C. Skins were washed and then secondary antibodies were added to PBT2. Skins were mounted in Prolong Gold or VectaShield. The following primaries were used: rabbit anti-Sox9 (1:1000, Millipore # AB5535), rat anti-E-Cadherin (1:1000, Invitrogen # 14-3249-82, clone DECMA-1), guinea pig anti-Celsr1 (1:1000, D. Devenport), chicken anti-GFP (1:2000, Abcam, ab6556), rabbit anti-RFP (1:200, Rockland 600-4010379). Appropriate Alexa Fluor-555, or 647 (Invitrogen) were used at 1:2000. Hoechst (Invitrogen #H1399) was used at 1 µg/mL with secondary. Z-stack images were acquired on an inverted Nikon A1R or A1R-Si confocal microscope controlled by NIS Elements software using a Plan Apo 60/1.4NA, Plan Apo 100/1.45NA oil immersion objective.

### Live Imaging

Live imaging was performed as previously described (Cetera et al., 2018). A 1% agarose gel with F-media supplemented with 10% fetal bovine serum was used for culture. Back skins were then dissected from E15.5 embryos and placed on the agar pad. The gel was sandwiched on a 35mm lummox dish (Starsted, product number: 94.6077.331) and imaged with an inverted Nikon inverted CSU-W1 SoRa Spinning Disk with Perfect Focus using a Plan Apo 20/0.75 NA air objective with a 2.8x zoom lens added to the light path. Explants were kept in a humid chamber with 5% CO2 at 37°C over the course of imaging. Timepoints were acquired every 20 minutes with a 2 µm step size for a total of 13-15 planes for at least 14 hours. Images were processed using NIS Elements and ImageJ. The MultiStackReg ImageJ plugin (Thévenaz et al., 1998) was used to correct for x-y drift using the dermis as a reference point. Positions were rotated using the average angle of guard hairs surrounding the placode of interest, to ensure the anterior was to the left. A single z-plane movie was created by following either the apical-most or basal-most surface of the placode through time.

### Inhibitor Treatment and Quantification

E15.5 dorsal skin explants were dissected in PBS and mounted on a 13 mm Whatman Nucleopore membrane with 8.0µM pore size. Explants were floated on F-media containing 10% fetal bovine serum. 10, 25, or 50 µM NSC 23766 in water (Rac1 inhibitor, Tocris Bioscience, NSC 23766, Cat:2161/10) was added to the F-media. Equivalent volume of water was used as control. Explants were cultured for 22 hours in 5% CO2 at 37°C and then fixed and stained as described in “Whole-mount immunostaining”. Hair placode polarity was quantified by a custom MATLAB script. Briefly, hair follicles and placodes were identified by thresholding on Sox9 channel intensity. A bounding box was then drawn around each placode. Polarity was calculated as the ratio of average Sox9 intensity on the posterior half of the bounding box and the anterior half.

### Image Segmentation and Polarity Analysis

Segmentation of hair placode cells of fixed and live images was performed using Cell Pose (Pachitariu and Stringer, 2022) with the membrane-tdTomato, membrane-eGFP, or E-cadherin channels. Segmentation masks were hand corrected in ImageJ using the Tissue Analyzer V2 plugin (Aigouy et al., 2016).

Polarity analysis was performed using Tissue Analyzer V2 in ImageJ (Aigouy et al., 2016). Tissue Analyzer was used to calculate the axis and magnitude of polarity of Celsr1, E-cadherin, Myosin IIB-GFP, and Fz6-3xGFP. Circular histograms plotting the data were generated using MATLAB (The MathWorks Inc., 2022), with average polarity magnitude indicating the angle and strength of polarity (Aw et al., 2016).

### Junction Analysis

Junction analysis was performed using ImageJ. Junctions of interest from time-lapse images were identified as either stable or dynamic using the membrane-tdTomato channel. Stable junctions were defined as junctions where length remained approximately consistent and was not associated with a neighbor exchange event for at least six consecutive time points (120 minutes). Dynamic junctions were defined as junctions associated with junction shrinkage to a vertex or a complete neighbor exchange. Junctions were then tracked for at least six frames (120 minutes) by drawing a rectangular ROI around the junction at each time point. Mean fluorescence intensity of either Fz6-3xGFP or Myosin IIB-GFP and junction length were measured for each time point. Measurements were then further processed using R 4.2.2 (R Core Team, 2022). For each dynamic junction, time points were binned into shrinking or growing by calculating the difference in junction length between a given timepoint and the previous one. Mean intensity values for each junction and timepoint were then normalized by the mean intensity of stable junctions from the corresponding movie. Statistical significance was tested using the unpaired Mann-Whitney U-test. Plot was generated using *ggplot2* (Wickham, 2016).

### Apical-Basal Tracking

From 3D time-lapse images (13-15 z-planes) of a placode from a *Fz6-3xGFP; mTmG* embryo, the apical and basal most planes of the hair placode were identified for each timepoint. These planes were then made into two time series, following the apical or basal surface until the basal-most surface was no longer in view. Each timepoint was then segmented based on membrane-tdTomato using CellPose (Pachitariu and Stringer, 2022) and hand corrected using Tissue Analyzer V2 (Aigouy et al., 2016). A custom MATLAB (The MathWorks Inc., 2022) applet was designed to allow for matching cells in both movies by hand. Inner cells were defined as cells whose centroid is within 15 µm of placode center. Outer cells were defined as cells whose centroid is greater than 15 µm of placode center (Leybova et al., 2024). Basal-Apical Offset over time plots were generated using MATLAB.

### 3D Fixed Reconstruction

3D reconstructions of the hair placode were generated as previous described (Leybova et al., 2024). 2D masks were generated using CellPose (Pachitariu and Stringer, 2022) from membrane-tdTomato or stained E-cadherin and hand-corrected for each frame in the z-stack (z-step of 0.25 µm or 0.5) using Tissue Analyzer V2 (Aigouy et al., 2016). The cells were first identified in each of the frames using the MATLAB (The MathWorks Inc., 2022) function *bwlabeln*. The regionprops function was then used to obtain the centroid of the cells in each z-slice. We implemented a distance-based heuristic algorithm to connect the centroids of the cells across the z-slices and create the 3D track of each cell. These centroid-connected cells were color-coded with the same color across frames, labeled and viewed in MATLAB’s *sliceViewer* for manual verification and correction. To produce a smooth 3D reconstruction of the cells and the placode, we added 5 (if z-step was 0.25 µm) or 10 (if z-step was 0.5 µm) interpolated slices between each consecutive real slice of the z-stack. The interpolated slices smoothly bridge the gap between the shape changes of each cell as they are tracked from frame to frame. The real data and interpolated slices were merged, generating a 3D matrix. The 3D view was constructed by plotting the isosurface of each cell. Each cell was assigned to a rainbow color based on distance from the centroid of the cell to the center of the placode/IFE region. Basal-Apical Offset for each cell was calculated as the x-component of the centroid of the apical minus that of the basal surface.

### Continuum Mechanics Model

Simulations were performed in a custom Python code (Python 3.10) using the Dedalus package to solve partial differential equations (Burns et al., 2020). Force field *F*_*x*_, and solutions for *ϕ* (density of Oldroyd-B polymers) and *u* (velocity field) are output from simulation every 10 iterations. Cell outlines consist of material points that track fluid flow. These points are moved every iteration by the interpolated velocity (Stein, 2022) using the second-order Adams-Bashforth method. Coordinates of tracked points are output every 10 iterations.

Calculations of metrics and vorticity were performed once the central cell has reached *x* = −6, as this corresponds approximately to the amount of deformation present in the placode when metrics were calculated experimentally. Basal-Apical Offset, Apical Shear, and Apical:Basal Shear Ratio were calculated using a custom MATLAB script (The MathWorks Inc., 2022). Vorticity (***∇*** × ***u***) of apical and basal surface velocities were calculated as the curl of the velocity field in Python and averaged over a time interval of *t* = 0.5 in a MATLAB script. Plots of cell outlines, phase diagrams for metrics, and density plots showing simulated fields (*F*_*x*_, *ϕ, u*, ***∇*** × *u*) are generated using custom Mathematica scripts (Wolfram Research Inc., 2025). Color map for velocity vector direction uses a perceptually uniform color scheme (Kovesi, 2015). See Supplemental Methods for full explanation of the mathematical model.

## Supplemental Movie Legends

**Movie S1. Live imaging of hair placode polarization in *Fz6-3xGFP; mTmG* embryo**. Time-lapse imaging of hair placode polarization from *E*15.5 embryo co-expressing membrane-tdTomato (inverted grayscale; top) and Fz6-3xGFP (bottom). 10.33 hrs/20 min interval. Scale bar: 10 µm.

**Movie S2. Live imaging of hair placode polarization in *Myosin IIB-GFP; mTmG* embryo**. Time-lapse imaging of hair placode polarization from *E*15.5 embryo co-expressing membrane-tdTomato (inverted grayscale; top) and Myosin IIB-GFP (bottom). Note the bright signal on the left of the placode is the myosin IIB within the dermal condensate. 11.33 hrs/20 min interval. Scale bar: 10 µm.

**Movie S3. 3D reconstruction of wild-type hair placode**. 3D rendering generated from fixed confocal images of hair placode from *E*15.5 *K14-Cre; mTmG* embryo. Cells are colored based on their distance from the center of placode (see Fig. 2A for color bar). Scale bars: 10 µm.

**Movie S4. 3D reconstruction of selected wild-type placode cells**. Selected examples of 3D reconstructed inner cells (red and orange, <15 µm from placode center) and outer cells (yellow and green, >15 µm from placode center). Cells are shown at their positions relative to the placode center. Apical is up. Scale bars: 5 µm.

**Movie S5. Live imaging of apical- and basal-most planes in polarizing hair placode**. Time-lapse imaging of hair placode polarization from *E*15.5 embryo expressing membrane-tdTomato (mT) following the apical-most (top) and basal-most (bottom) planes. Note the faint signal surrounding placode cells is mT expression in dermal fibroblasts. 10.33 hrs/20 min interval. Scale bar: 10 µm.

**Movie S6. 3D reconstruction of *Celsr1***^***KO/KO***^**; *mTmG* hair placode**. 3D rendering generated from fixed confocal images of hair placode from *E*15.5 *Celsr1*^*KO/KO*^; *mTmG* embryo. Cells are colored based on their distance from center of placode (see Fig. 3A for color bar). Scale bars: 10 µm.

**Movie S7. 3D reconstruction of selected *Celsr1***^***KO/KO***^**; *mTmG* placode cells**. Selected examples of 3D reconstructed inner cells (red and orange, <15 µm from placode center) and outer cells (yellow and green, >15 µm from placode center) from *Celsr1*^*KO/KO*^; *mTmG* embryo. Cells are shown at their positions relative to the placode center. Apical is up. Scale bars: 5 µm.

**Movie S8. Evolution of tracked material points of inner and outer cells in simulation with uniform stiffness and weak apical adhesion**. Cell outlines at basal (left) and apical (center-left) surface of placode viscoelastic slab region *S*. Anterior-directed crawling force magnitude shown in shades of blue at basal surface (left, see Fig. 5D for color bar). Basal and apical surfaces shown in 3D view (center-right). 3D wireframe of inner cells shown in 3D view (right). Simulation parameters: *F*_*x*_ = 64. *H*_2_ = 0.5. *η*_*A*_ = 0.05. *ϕ* = 9 uniform. Anterior is to the left. Simulation ends once center cell reaches *x* = −6, at *T* = 38.

**Movie S9. Evolution of tracked material points of inner and outer cell surfaces for different simulation conditions of apical adhesion and tissue stiffness**. Cell outlines at apical (top panels) and basal (bottom panels) surface of placode viscoelastic slab region *S*. Crawling forces shown in shades of blue at basal surface (bottom panels, see Fig. 5D for color bar). This is shown for three simulation conditions, uniform stiffness and weak apical adhesion (left), uniform stiffness and strong apical adhesion (middle), and non-uniform tissue stiffness and weak apical adhesion (right). Simulation parameters: for all three, *F*_*x*_ = 64. Left: *H*_2_ = 0.5, *η*_*A*_ = 0.05, *ϕ* = 9 uniform. Middle: *H*_2_ = 0.05, *η*_*A*_ = 2.0, *ϕ* = 9 uniform. Right: *H*_2_ = 0.5, *η*_*A*_ = 0.05, *ϕ*_*IFE*_ = 18, *ϕ*_*Outer*_ = 1, *ϕ*_*Inner*_ = 9 (see Fig. 6G left). Each simulation ends once center cell reaches *x* = −6. Note that simulations with weak apical adhesion (left and right) reach *x* = −6 at approximately at the same time (*T* = 38 and *T* = 36, respectively), whereas strong apical adhesion (middle) reaches this point later *T* = 46.75.

**Movie S10. Surface velocity fields of apical and basal forces for different simulation conditions of apical adhesion and tissue stiffness**. Velocity fields of the apical (top panels) and basal (bottom panels) surfaces of the model slab shown for three simulation conditions: uniform stiffness and weak apical adhesion (left), uniform stiffness and strong apical adhesion (middle), and non-uniform tissue stiffness and weak apical adhesion (right). Simulation parameters: for all three, *F*_*x*_ = 64. Left: *H*_2_ = 0.5, *η*_*A*_ = 0.05, *ϕ* = 9 uniform. Middle: *H*_2_ = 0.05, *η*_*A*_ = 2.0, *ϕ* = 9 uniform. Right: *H*_2_ = 0.5, *η*_*A*_ = 0.05, *ϕ*_*IFE*_ = 18, *ϕ*_*Outer*_ = 1, *ϕ*_*Inner*_ = 9 (see Fig. 6G left). Note that simulations with weak apical adhesion (left and right) reach *x* = −6 at approximately the same time (*T* = 38 and *T* = 36, respectively), strong apical adhesion (middle) reaches this point later *T* = 46.75.

**Movie S11. Evolution of average vorticity for different simulation conditions of apical adhesion and tissue stiffness**. Average vorticity fields of the apical surface velocity. Vector length is proportional to velocity magnitude and colored based on angle from -180° to 180°. Average calculated over time windows of *t* = 0.5 (see Methods). This is shown for three simulation conditions, uniform stiffness and weak apical adhesion (left), uniform stiffness and strong apical adhesion (middle), and non-uniform tissue stiffness and weak apical adhesion (right). Simulation parameters: for all three, *F*_*x*_ = 64. Left: *H*_2_ = 0.5, *η*_*A*_ = 0.05, *ϕ* = 9 uniform. Middle: *H*_2_ = 0.05, *η*_*A*_ = 2.0, *ϕ* = 9 uniform. Right: *H*_2_ = 0.5, *η*_*A*_ = 0.05, *ϕ*_*IFE*_ = 18, *ϕ*_*Outer*_ = 1, *ϕ*_*Inner*_ = 9 (see Fig. 6G left). Note that simulations with weak apical adhesion (left and right) reach *x* = −6 at approximately the same time (*T* = 38 and *T* = 36, respectively), strong apical adhesion (middle) reaches this point later *T* = 46.75.

**Figure S1.**
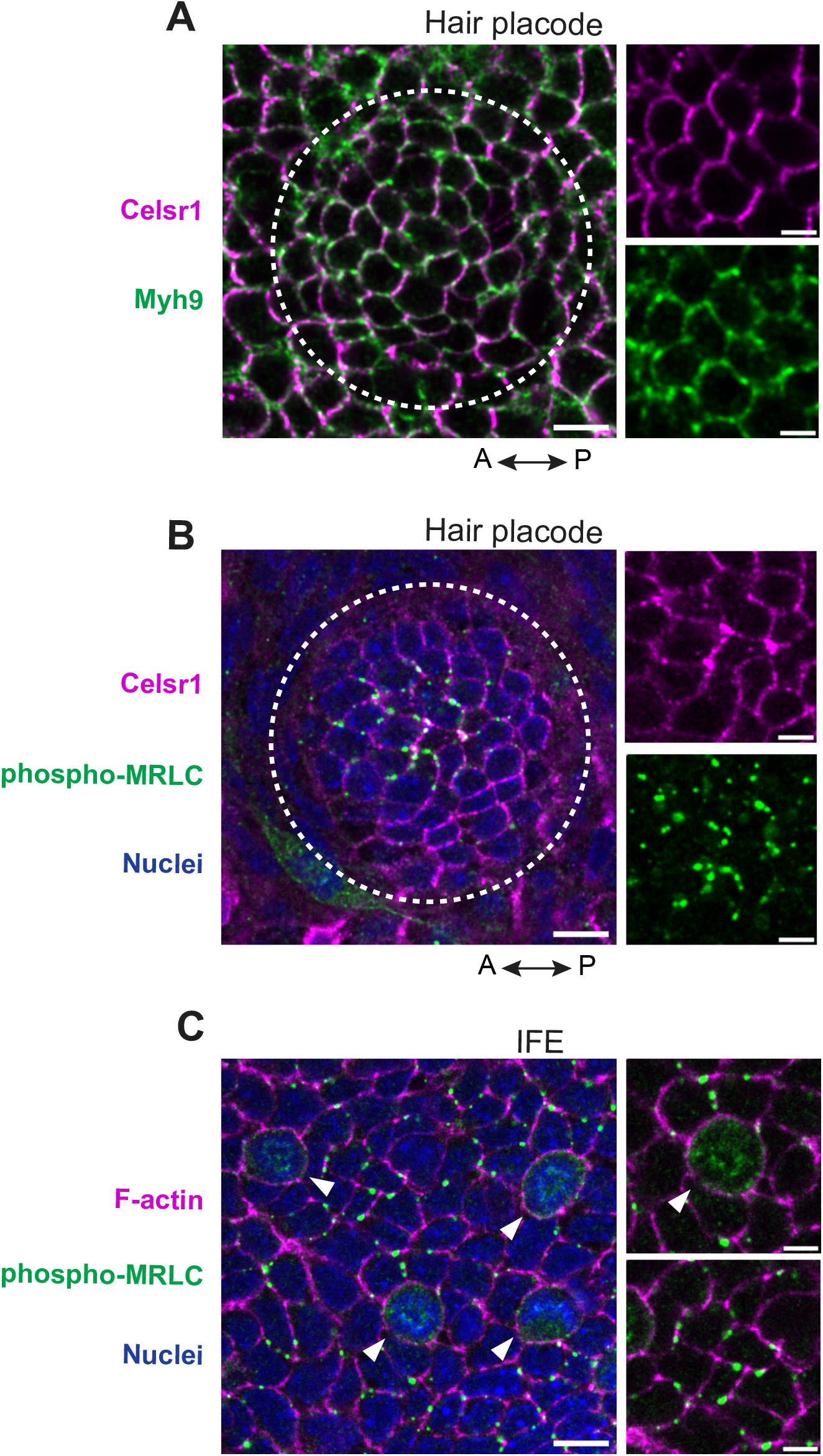
Localization of total and phosphorylated non-muscle myosin in hair placodes at polarization stages. A) Representative hair placode from *E*15.5 whole mount epidermis immunolabeled with Celsr1 (magenta) and non-muscle myosin heavy chain 9 (MYH9, green) antibodies. Note the lack of colocalization between Celsr1 and MYH9 at epithelial junctions. Anterior to left. Scale bars: 10 μm, insets 5 μm. B) Representative hair placode from *E*15.5 whole mount epidermis immunolabeled with Celsr1 (magenta) and phosphorylated myosin regulatory light chain (p-MRLC, green) antibodies. p-MRLC labels foci located along epithelial junctions mainly in the center (inner cells) of the placode. However, these foci do not strongly correlate with regions of Celsr1 intensity. Anterior to left. Scale bars: 10 μm, insets 5 μm. C) Representative image of the interfollicular epidermis (IFE) from *E*15.5 whole mount epidermis labeled with phalloidin (F-actin, magenta) and phosphorylated myosin regulatory light chain (p-MRLC, green) antibodies. p-MRLC labels foci located along epithelial junctions of the IFE and as expected and is elevated at the cortex and cytoplasm of cells in mitosis (arrowheads). Scale bars: 10 μm, insets 5 μm.

**Figure S2.**
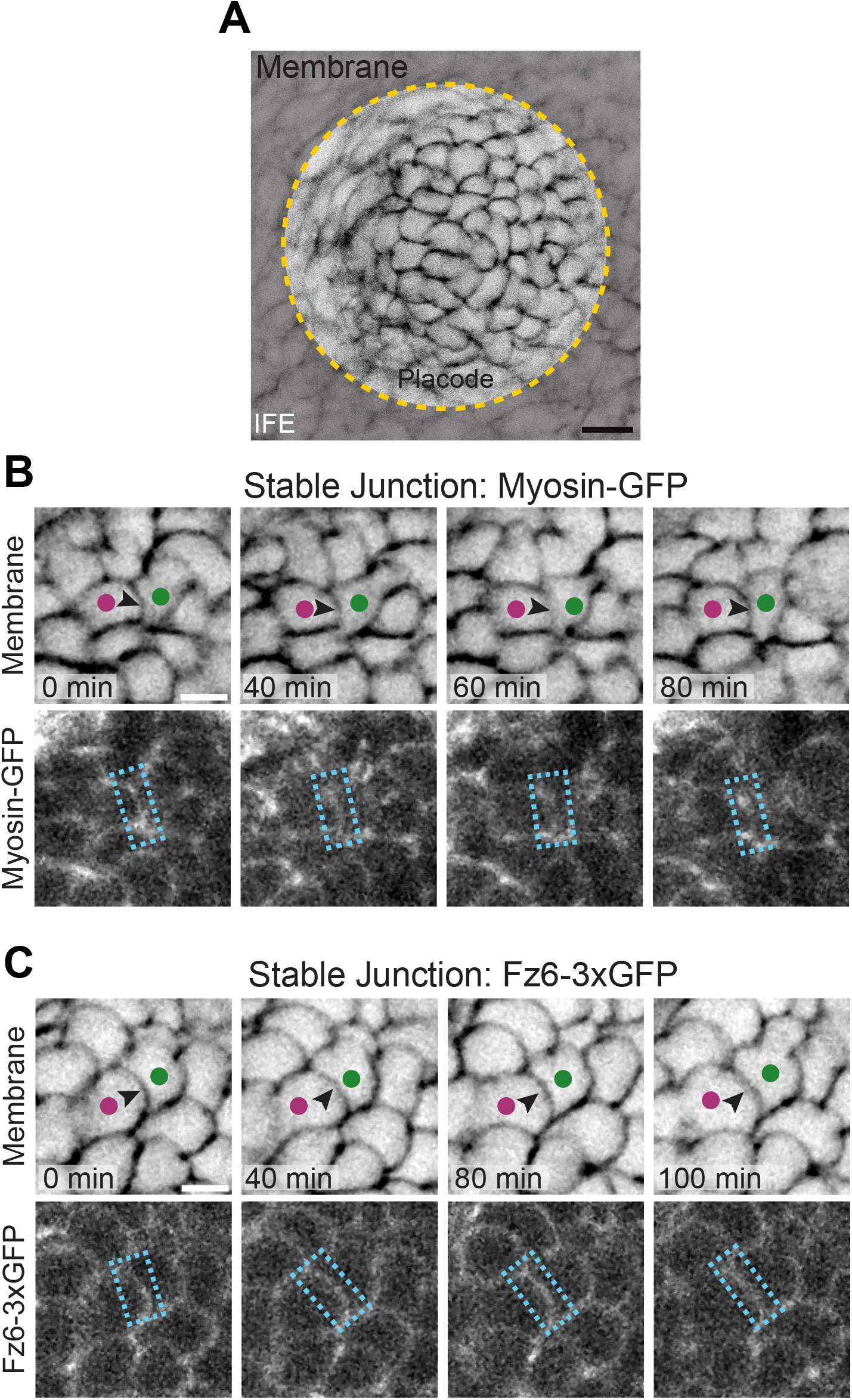
Myosin and Fz6 localization through time at stable junctions during placode polarization. (A) Image depicting representative placode from *E*15.5 skin explants expressing Myosin IIB-GFP and membrane-tdTomato. Time lapse images in Fig 1E,G and S2 are taken from the center of a developing hair placode during polarization. Scale bar: 10μm. B) Representative time-lapse images from *E*15.5 skin explants co-expressing membrane-tdTomato (upper panels) and Myosin IIB-GFP (lower panels). A stable junction (arrowhead) between two cells that retain their positions is shown (stable – blue boxes). Scale bar: 5 μm. C) Representative time-lapse images from *E*15.5 skin explants co-expressing membrane-tdTomato (upper panels) and Fz6-3xGFP (lower panels). A stable junction is shown between two cells that retain their positions through time (stable – blue boxes). Scale bar: 5 μm.

**Figure S3.**
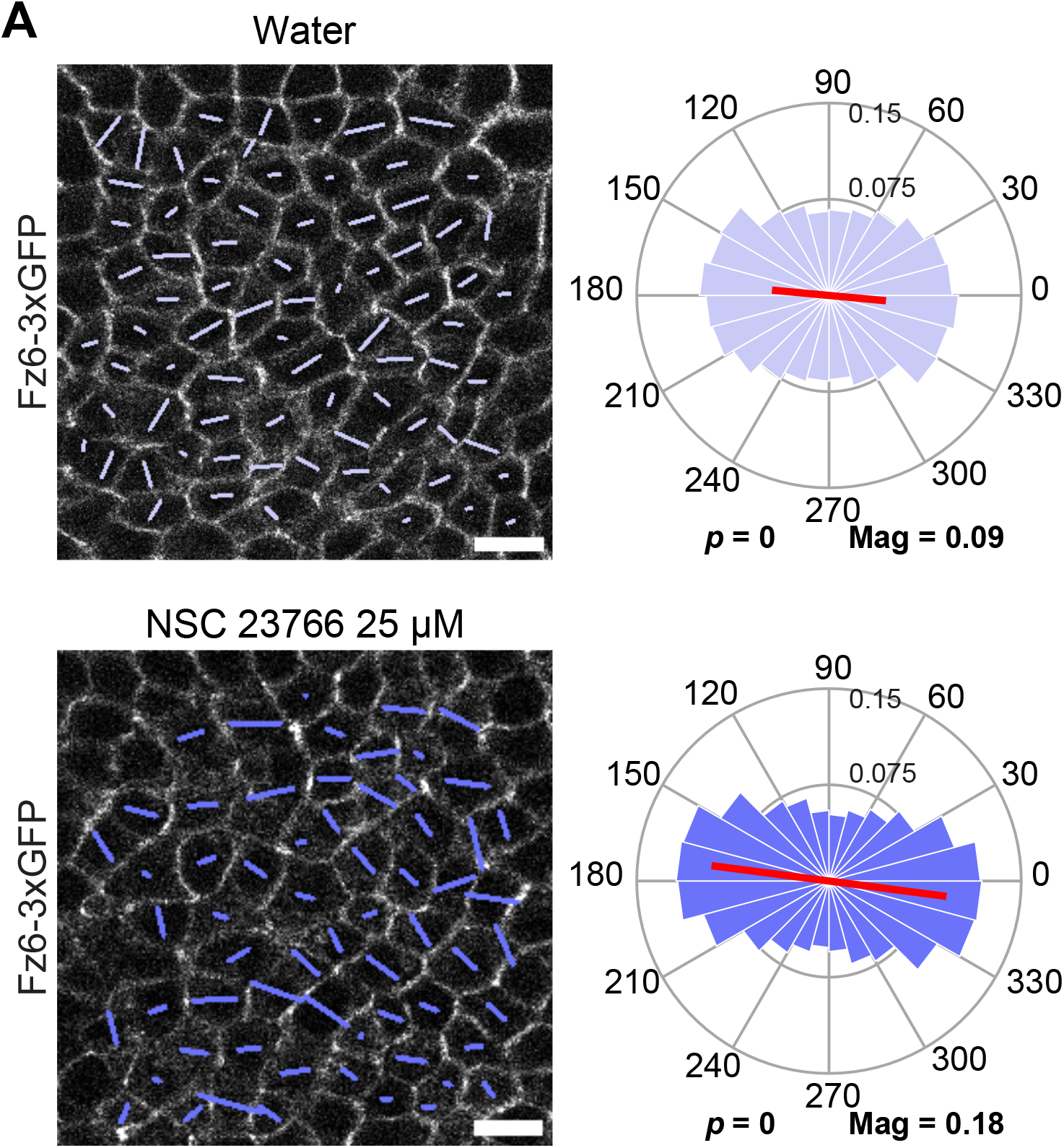
Fz6 remains planar polarized in embryonic skin explants cultured in Rac1 inhibitor. A) Representative planar views of hair placode from *E*15.5 embryo expressing Fz6-3xGFP cultured for 22 hours with water as a control (top panel) or 25μM NSC 23766 (bottom panel). Skin explants were immunolabeled with antibodies against Celsr1 and E-cadherin. Images are overlaid with polarity nematic quantifying polarization of Fz6-3xGFP (left panel). Quantification of polarity distributions are displayed below as circular histograms (right panel). Scale bar: 10 μm.

## Supplemental Computational Methods

### 1 Model Equations

The Oldroyd-B viscoelastic fluid model presented here heavily references the methods from [1], but with extensions to the model, presented below.

We model the developing hair placode and surrounding cells as a viscoelastic slab with surrounding fluid layers and hard, rigid walls. We represent the developing mouse hair placode plus the surrounding interfollicular epithelium (IFE) as an Oldroyd-B viscoelastic fluid occupying a slab-shaped region (region *S*). The placode cells plus the IFE make up the “epidermal basal layer” in scientific nomenclature. However, for this work, to avoid confusion and multiple uses of the word “basal”, we will term this collection of cells the “Placode Layer”. We model the Placode Layer as a flat slab because it is significantly larger in the *x* and *y* directions and than the *z* direction. Moreover, although the placode deforms in the *z* direction, it remains nearly flat during the specific stage of polarization that we are interested in, so we keep it flat in the model. Hence, to describe a single placode surrounded by IFE cells, we model a 3D slab that is periodic in the *x* and *y* directions with thickness *h* in the *z* direction.

Our length and timescales of interest are between 0.1 and 100 microns and 1 minute to 300 minutes, respectively. This covers the length and time scales of the adhesion interface between tissues, as well as that of multicellular rearrangements.

#### 1.1 Equations of motion for the epithelium and surrounding material

In the mouse hair placode, there is material above (apical to) and below (basal to) the Placode Layer. We model these as regions *A* and *B*, and we model the materials comprising these regions as Newtonian fluids. Region *A* represents the extracellular space between the Placode Layer and the suprabasal cells plus the basal-most (∼ 2*µ*m) portion of the suprabasal cells, a region that is more flexible than the rest.

Region *A* resides on the apical side of the Placode Layer, and we refer to it as the “apical fluid layer” in the model. Biologically, we refer to it as the “apical adhesion zone” or “apical adhesion interface”. Region *B* represents the extracellular space between the Placode Layer and the basement membrane plus the relatively flexible portion of the basement membrane (*lamina lucida*) that is adjacent to the Placode Layer. Region *B* resides on the basal side of the Placode Layer, and we refer to it as the “basal fluid layer” in the model. In the model, both regions *A* and *B* are bounded by hard walls. The hard wall on the apical side represents the remainder of the suprabasal cells, beyond the basal-most 2*µ*m, where they become stiffer. The hard wall on the basal side represents the *lamina densa* (the stiffer part) of the basement membrane. Both of these walls are assumed to be anchored, consistent with experimental observations, as microscopy data shows that these sections are immobile throughout polarization of the placode.

We set up our coordinate system so that region *B* corresponds to 0 < *z* < *H*, region *S* corresponds to −*h* < *z* < 0, and region *A* corresponds to −*h* −*H*_2_ < *z* < −*h*. This coordinate system therefore assigns height *H* to the basal fluid layer; it assigns height *h* to the Placode Layer, and it assigns height *H*_2_ to the apical fluid layer. Figure S4 shows regions *B, S*, and *A* in a schematic diagram along with the coordinate system we use to simulate the placode. Notably, we solve the system with the basal side of the placode geometrically above the apical side, contrary to the orientation in which biological data is often presented. This is an arbitrary choice that does not affect calculations. However, for clarity and for correspondence with biological conventions, we plot simulations with *z* → − *z* in both the main text and the Supplemental Movies. Figure 1 of the main text shows the system orientation as the simulations are plotted, with apical side up.

**Figure S4:**
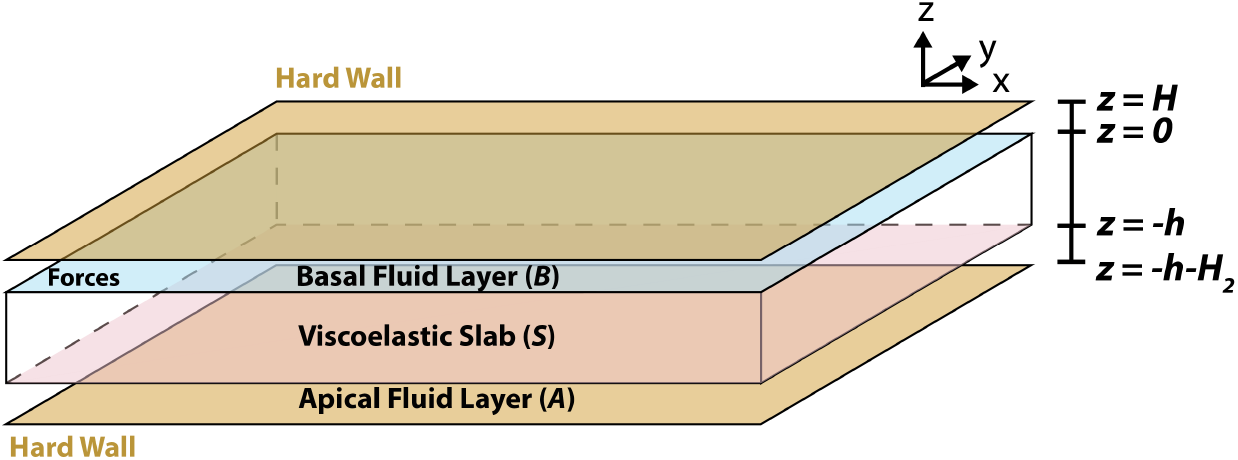
Schematic diagram of model epithelium in the slab geometry with definitions of regions *B, S, A*, and the coordinate system. The basal fluid layer, Placode Layer epithelium, and apical fluid layer have thicknesses *H, h*, and *H*_2_, respectively.

Similarly to [1], we model stresses in regions *B, S*, and *A* of the system as:

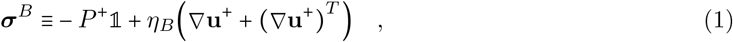

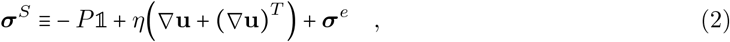

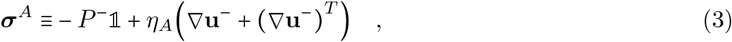

where **u, u**^±^ are the velocities, *P, P* ^±^ are the pressures, and *η, η*_*B,A*_ are the viscosities in each region. Here, ***σ***^*B,A*^ indicate Newtonian fluids and ***σ***^*e*^ indicates an extra viscoelastic stress in region *S*, which we describe below. The above expressions for the stresses, along with the requirements that forces are balanced:

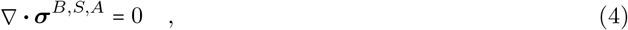

and that fluids are incompressible:

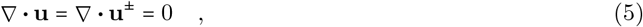

result in the equations of motions for **u, u**^±^ and *P, P* ^±^ which we summarize in dimensionless form at the end of this section.

We complete our description of the model by giving the equations of motion for ***σ***^*e*^, the extra stress coming from the Oldroyd-B description of the tissue. Heuristically, an Oldroyd-B fluid describes a viscous fluid with Oldroyd-B polymers (also called Oldryod-B particles) embedded in the fluid. These polymers are modeled as balls connected by springs. As the fluid flows, the polymer springs stretch and relax. When this model (often called the “kinetic theory”) is taken to the continuum limit in a mathematical procedure called coarse-graining, these Oldroyd-B particles contribute an associated stress to the system, called the extra viscoelastic stress (***σ***^*e*^). This stress contributes to the forces in the equations of motion alongside the viscous stress and the pressure. The quantity ***σ***^*e*^ in Eq. (2) is evolved by the equation:

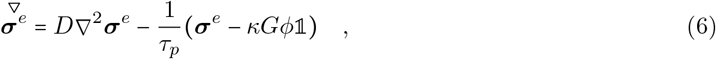

in which

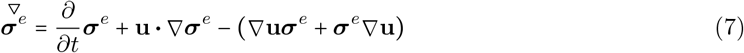

is defined as the upper-convected time derivative. Here, *τ*_*p*_ represents the relaxation time of Oldroyd-B polymers while *κ* and *G* are parameters associated with the kinetic theory of Oldroyd-B [2], and *D* is the center of mass diffusion of Oldroyd-B polymers. The quantity *ϕ* is a field representing the density of Oldroyd-B polymers whose evolution is explained below. Using the kinetic theory, if we assume that Oldroyd-B polymers cannot diffuse across boundaries of the tissue (conserving particle numbers), then this implies that the diffusive flux at the boundaries *z* = 0 and *z* = −*h* are both 0:

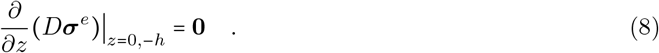

The field *ϕ* (**x**, *t*) in Eq. 6 represents the density of Oldroyd-B particles at position **x** and time *t*. It is evolved according to:

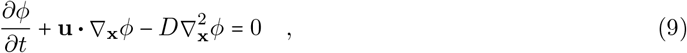

with zero diffusive flux boundary conditions given by

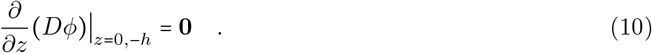

Section 1.7 provides the derivation of Eq. 9 from the kinetic theory. Equations 6 and 9 along with Eqns. 4 and 5 (and joined by definitions 1-3 and 7) form the complete set of equations for the system.

#### 1.2 Summary of non-dimensionalized equations

To non-dimensionalize Eqns. 1-9, we follow similar procedures as in [1] and rescale time, length, and force as:

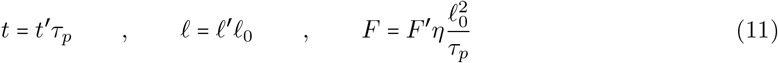

where *ℓ*_0_ is a characteristic length scale that we eventually set to the length of one hexagonal cell’s edge. We choose dimensional units to depend on the material properties of the system (the solvent viscosity *η* and polymer relaxation time *τ*_*p*_), with the exception of *ℓ*_0_ which is dependent on cell size. This choice of time, length, and force scale effectively sets *τ*_*p*_, *η*, and the edge length of a cell all to the value 1 when the equations are in dimensionless form. The details of non-dimensionalization are summarized below and presented in greater detail in the Supplement of [1]. We start with:

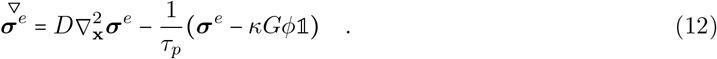

To rescale Eq. 12, we replace dimensional parameters and fields with dimensionless quantities multiplied by the correct dimensional unit. We use primes to indicate dimensionless quantities:

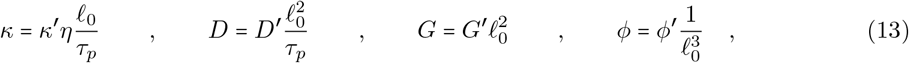

In anticipation of algebraic simplification, we rescale ***σ***^*e*^ with additional dimensionless numbers *κ*^′^, *G*^′^, and *ϕ*^′^:

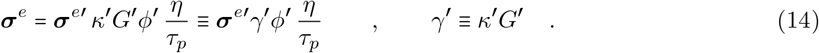

Note that *ϕ*^′^ and 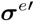 are dimensionless fields whereas other primed quantities are dimensionless numbers. The dimensionless derivatives are:

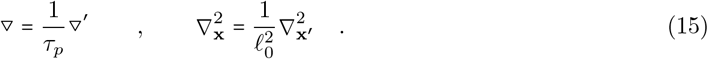

Substituting these into Eq. 12, we have:

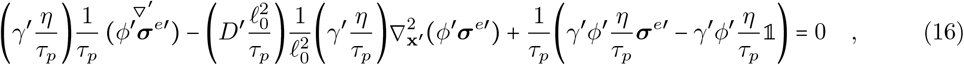

where we used that 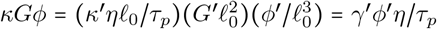. Eliminating constants, the above simplifies to:

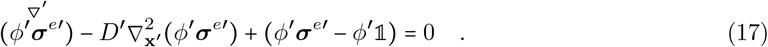

From here, denote 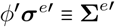, then the above becomes:

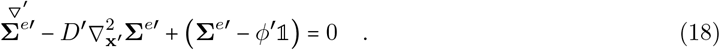

Let us now consider:

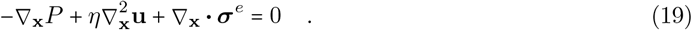

Expressing all fields as the multiplication of a non-dimensional field with a dimensional unit:

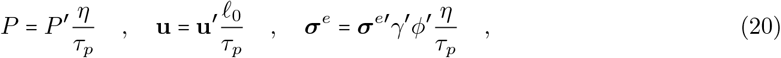

substituting these into Eq. 19, and recalling that *ϕ*^′^ is a field and has to stay behind the divergence, we have:

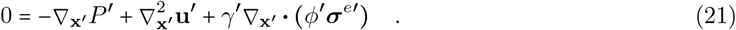

Using 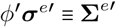, this becomes:

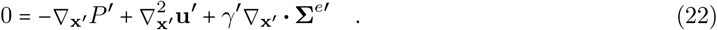

Finally, for *ϕ*, we have:

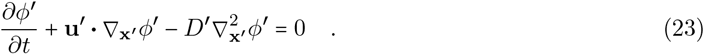

Equations 18, 22, and 23 are the non-dimensional equations for the viscoelastic Oldroyd-B fluid in region *S* of the system.

We summarize below the non-dimensional equations of motions for regions *B, S*, and *A*. We remove all primes for notational clarity:

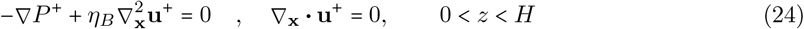

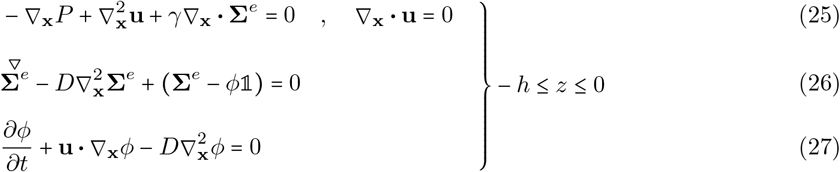

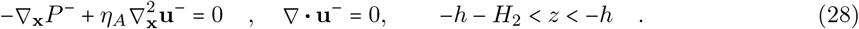

To obtain solutions to our model, we solve for the dimensionless fields **Σ**^*e*^ (instead of ***σ***^*e*^), *ϕ*, **u, u**^±^, *P*, and *P* ^±^.

#### 1.3 Boundary conditions between regions

To solve for all pressures *P, P* ^±^ and all velocities **u, u**^±^ in our system, we must specify velocity and/or stress boundary conditions on all surfaces. First, we assume a no-slip boundary condition on both hard walls at *z* = *H* and *z* = −*h* − *H*_2_,

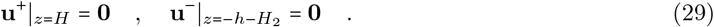

Additionally, as in [1], we assume that at *z* = 0 and *z* = −*h*, the interfaces between the slab and the fluid layers *A* and *B*, we have continuity of velocity in the *x* and *y* directions as well as a no-penetration zero-velocity condition in the *z* direction:

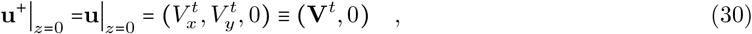

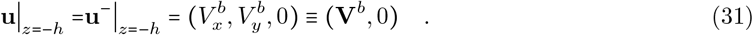

Here, using similar notation as [1], the planar *x*-*y* velocity at *z* = 0 is denoted 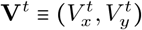, where the superscript denotes the “top” surface of the slab (basal side of the epithelium), and the planar velocity at *z* = −*h* is denoted 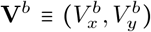 where the superscript denotes the “bottom” surface of the slab (apical side of the epithelium).

In order to incorporate active tangential or planar forces applied at the apical and basal side of the Placode Layer, we allow for active driving forces at the surfaces of the slab at *z* = 0 and *z* = −*h*. These active forces correspond to stress differences (often called stress jumps) across those surfaces, and they are reflected in the boundary conditions. Let **F**^*t*^ and **F**^*b*^ notate the tangential stresses (force per unit area) from active driving forces applied at the top (basal) and bottom (apical) surfaces, respectively.

Then the stress boundary conditions are:

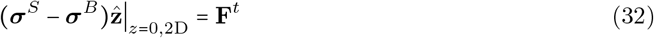

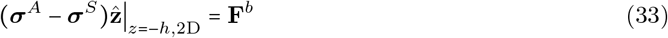

where the notation 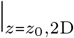 is defined to notate taking only the first 2 components of a vector and evalulating at *z*_0_, that is, 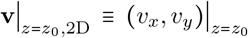 for an arbitrary 3D vector **v**= (*v*_*x*_, *v*_*y*_, *v*_*z*_). The *z*-directional stress at the top and bottom forces at the slab surfaces are determined by the non-penetration velocity boundary condition at *z* = 0, *h*.

#### 1.4 Solutions to the system and the Neumann-to-Dirichlet map

In order to solve the system for *P, P* ^±^, **u, u**^±^, we follow the methods in [1] and start by solving the Stokes system without Oldroyd-B. This means that we set **Σ**^*e*^ = **0** and eliminate Eqns. 26 and 27. The analytical expressions for the solutions to this Stokes-only system are detailed in Section 1.6. Central to solving this Stokes-only system is to perform the 2D Fourier Transform of Eqns. 24, 25, and 28 in the *x* and *y* directions. The 2D Fourier Transform of an arbitrary scalar function *f* is defined as:

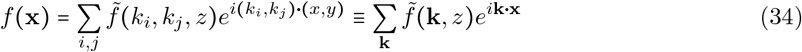

where **k** takes on values:

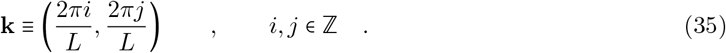

This transformation allows for solutions of the Stokes-only system that provide analytical expressions to *P, P* ^±^, **u, u**^±^ and their Fourier Transforms 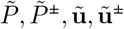, which are used in the following expressions. Specifically, the Fourier transformed expressions 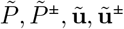 depend linearly on 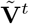 and 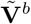 which are the Fourier Transforms of the boundary surface velocities **V**^*t*^ and **V**^*b*^. See Section 1.6 for full mathematical expressions. From expressions 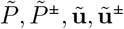, we compute the shear stresses at both *z* = 0 and *z* = −*h* coming from the fluid layers *A* and *B* and coming from the slab *S*. These shear stresses are defined as the projection of the stress tensor in these regions on the *z* direction at *z* = 0, −*h*. Notably, their expressions in Fourier form also depend linearly on 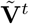 and 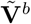:

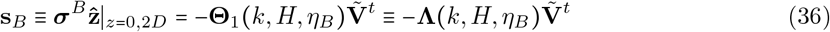

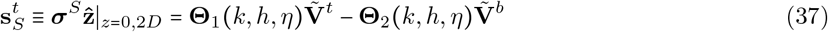

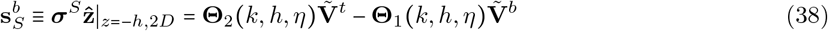

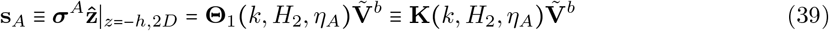

The expressions **Θ**_1_ and **Θ**_2_ are 2 × 2 matrices that multiply the *x* and *y* components of 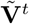 and 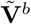; they depend on the magnitude *k* of the wavevector **k** as well as the region geometry (*H, h*, or *H*_2_) and their viscosities (*η*_*B*_, *η*, or *η*_*A*_). Here, we defined **Λ** as **Θ**_1_ evaluated with the arguments (*k, H, η*_*B*_), and we defined **K** as **Θ**_1_ evaluated with the arguments (*k, H*_2_, *η*_*A*_). These definitions are simply to enable us to use **Θ**_1_ and **Θ**_2_ as shorthand for **Θ**_1,2_ (*k, h, η*), evaluated with the arguments (*k, h, η*) by default.

The expressions for **Θ**_1,2_ are given in the Supplement for [1], copied below to aid the reader:

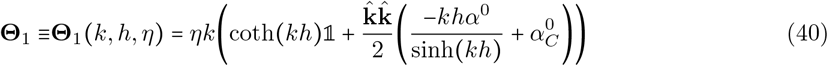

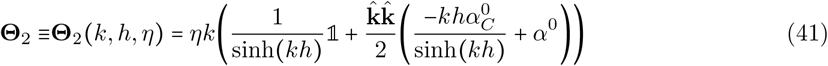

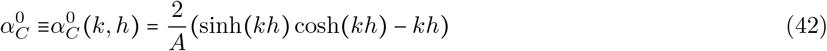

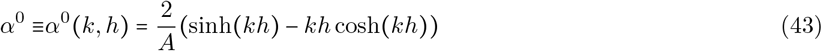

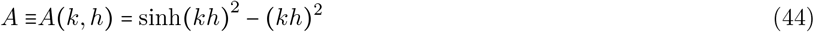

Note that all expressions so far are linearly dependent on 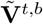, but 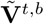 have yet to be specified. The quantities that *are* specified are the active forcing 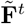 and 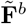 (the Fourier transforms of **F**^*t*^ and **F**^*b*^) that are given by the forcing that we specify in the problem. Hence the missing step is to express 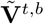 in terms of 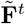 and 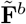 to complete the solution for the system. This is done through the Neumann-to-Dirichlet map which defines the translation from stress or force (Neumann) boundary conditions that specify 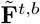 to velocity (Dirichlet) boundary conditions 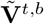 that then enter the expressions of interest. Writing the stress boundary conditions in Eqns. 32 and 33 in Fourier form, we have

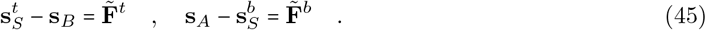

The above combined with the expressions for the shear stresses in Eqns.36-39 give rise to the equations:

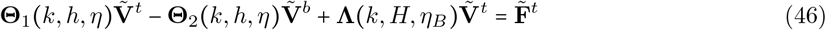

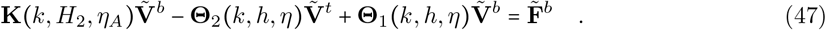

The above expressions are inverted so as to express 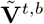 in terms of 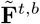. Finally, defining

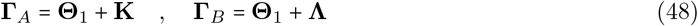

and using the shorthand that **Θ**_1_ and **Θ**_2_ refer to **Θ**_1,2_ evaluated with the arguments (*k, h, η*) while **Λ** ≡ **Θ**_1_(*k, H, η*_*B*_) and **K** ≡ **Θ**_1_(*k, H*_2_, *η*_*A*_), we have the Neumann-to-Dirichlet maps:

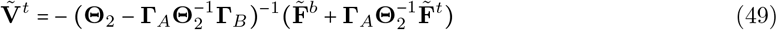

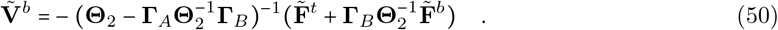

From the above, given any active forces 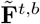 that are applied to either the top or bottom of the slab (the Placode Layer in our model tissue), we can evaluate 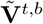 and proceed to express all field solutions *P, P* ^±^, **u, u**^±^ in terms of 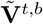.

#### 1.5 Incorporating the Oldroyd-B extra stress

The previous section gave the formulation of how to determine all pressures and velocities in the system (analytically) given the active force in the absence of the viscoelastic extra stress that comes from the Oldroyd-B polymers. Here, we re-incorporate the effects of the Oldroyd-B polymers numerically. The details of how to precisely formulate the incorporation of the Oldroyd-B polymers are given in Section III of the Supplement to [1]. Below, we simply provide the computation formula in the presence of Oldroyd-B. First, solve the numerical problem:

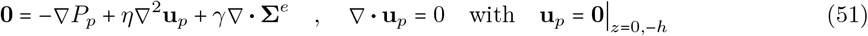

for fields *P*_*p*_ and **u**_*p*_ in region *S*. Note that **Σ**^*e*^ is simply a 3 × 3 numerical field at every time point. Also note that here, we impose that the velocities are identically **0** at the boundaries *z* = 0, −*h*, which makes this a straightforward problem with Dirichlet boundary conditions to be solved numerically. Next, evaluate the 3D Newtonian stress tensor ***σ***_*N*_ (*P*_*p*_, **u**_*p*_, *η*) = − *P*_*p*_𝟙 + *η*(∇ **u**_*p*_ + (∇**u**_*p*_) ^*T*^) that depends on the numerical solutions *P*_*p*_, **u**_*p*_ that were found. Finally, compute:

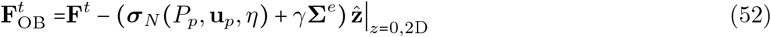

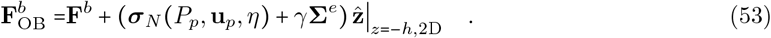

Taking the Fourier transform of the above as 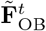 and 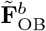, we substitute them into Eqns. 49 and 50 in place of 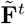 and 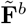, respectively. The resulting expressions then give the surface velocities in terms of the active forces in the presence of Oldroyd-B particles:

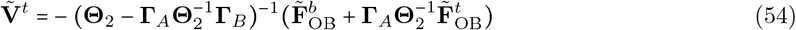

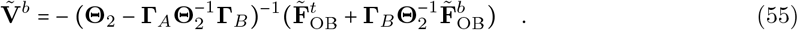

The above 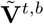 are then used to compute *P, P* ^±^, **u, u**^±^, as before. Finally, the full pressure and velocity solutions in region *S* is amended to:

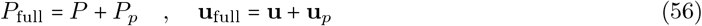

The effect of adding the Oldroyd-B polymers to region *S* is that the bulk solutions in the slab are amended from the Stokes-only solution to include the effects of the polymers; additionally, extra stress from the polymers are induced at the surface and added to active stresses **F**^*t*^ and **F**^*b*^ to form 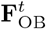 and 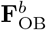, which then affects surface velocities **V**^*t*^ and **V**^*b*^.

#### 1.6 Solutions to Bulk Equations

We here present the homogeneous solutions for velocity and pressure (setting ***σ***^*e*^ = **0**) to Eqns. 4 with boundary conditions in Eqns. 30 and 31. We mention here that the full solutions in regions *B* and *A* are simply the homogeneous solutions, while the solution in region *S* requires the addition of *P*_*p*_ and **u**_*p*_ to the homogeneous solutions, as noted in Eq. 56. As previously mentioned, *P*_*p*_ and **u**_*p*_ are numerical solutions to Eq. 51. Hence, the homogeneous solutions to all regions presented in the following will complete the calculation of bulk solutions. The solutions here are similar to the ones given in Section IIA of the Supplement of [1], with some changes due to the differences in geometry.

##### 1.6.1 Velocity and pressure solutions in region *S*

Recall that the velocity in this region is denoted **u** = (**v**, *w*), and the pressure denoted *P* . Let 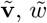, and 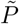 define the Fourier Transformed version of these fields via Eq. 34. To simplify expressions, we recall definitions in Eqns. 42-44 and additionally define:

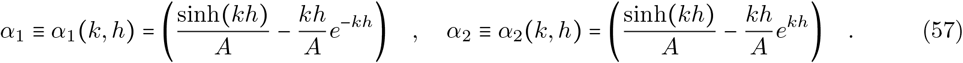

This gives that 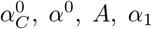, and *α*_2_ are all parameter-free functions of (*k, h*). We have for 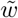 in region *S*:

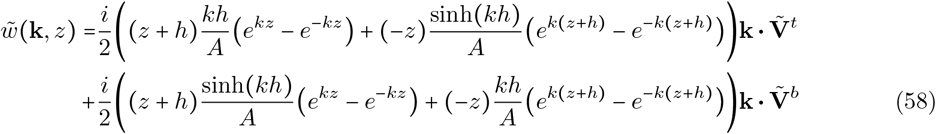

and for 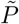 in region *S*:

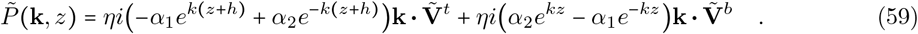

Finally, the solution for 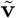, the *x, y* velocity, in region *S* is:

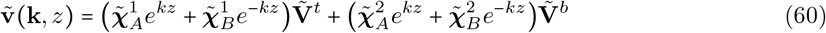

where 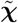’s are matrices multiplying the 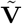 vectors given by:

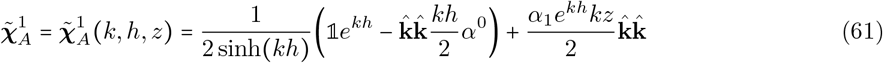

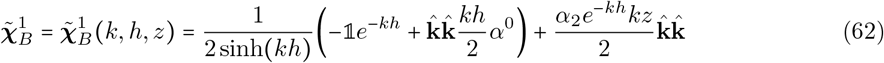

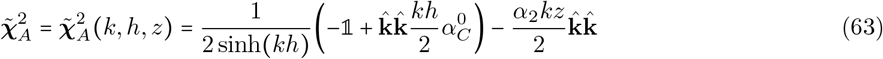

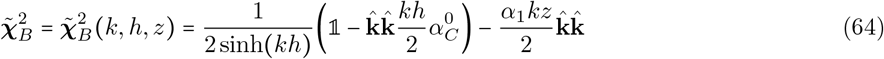

##### 1.6.2 Velocity and pressure solutions in region *B*

We note that the solutions here are nearly identical to the ones listed in Section IIA(2) of the Supplement in [1], with the vicosity of this region *η*_*B*_. The solution for 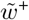 in region *B* is:

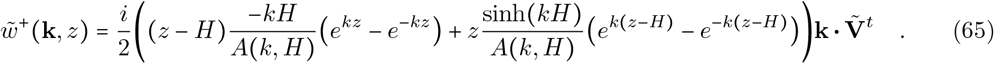

For 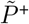 in region *B*:

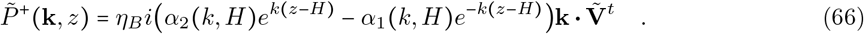

For 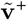 in region *B*:

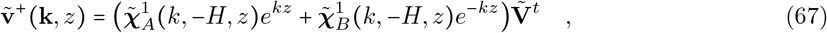

where

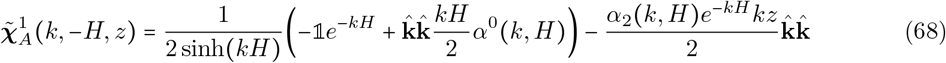

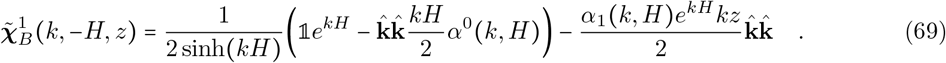

##### 1.6.2 Velocity and pressure solutions in region *A*

We note that the solutions here can be adapted from the solutions in Eqns. 58, 59, and 60 by shifting them. First, we make the replacement that *h* → *H*_2_, as the thickness of region *A* is *H*_2_. Next we replace the argument *z* → *z* + *h*, as the top of region *A* at *z* = −*h* should be mapped to 0 in the [− *h*, 0] geometry. We also note that the “top” velocity 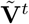 in Eqns. 58, 59, and 60 should be replaced by 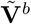 as this is now the velocity at the top of region *A*, and the “bottom” velocity in Eqns. 58, 59, and 60 should be replaced by **0**, as this should denote the velocity at the wall. Lastly, we note that the vicosity of this region *η*_*A*_. We have for 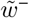 in region *A*:

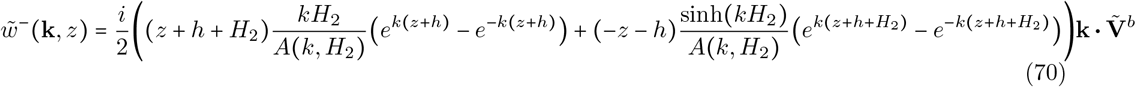

and for 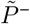 in region *A*:

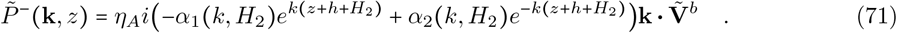

Finally, the solution for 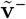, the *x, y* velocity in region *A*, is:

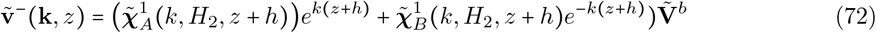

with

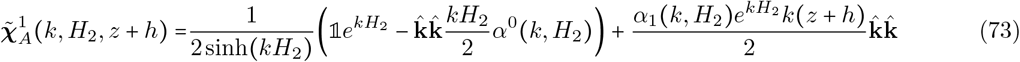

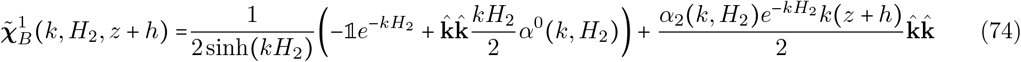

#### 1.7 Derivation of equation of motion of density *ϕ*

From the kinetic theory of Oldroyd-B [2], the scalar field *ϕ* (**x**) is defined as:

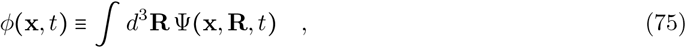

where Ψ (**x, R**, *t*) is the density of polymers at position **x** with orientation **R**. To derive the equation of motion for *ϕ*, we take the derivative of *ϕ* with respect to time:

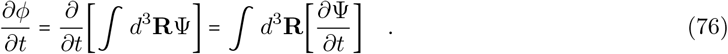

From the Smoluchowski equation in [2], we have

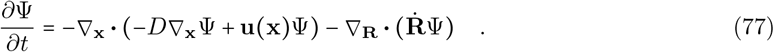

Hence, substituting Eq. (77) into Eq. (76) and pulling the ∇_**x**_ through, we have:

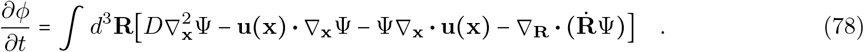

By the divergence theorem, we have that 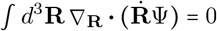. Hence the above simplifies to:

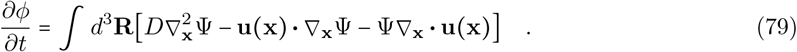

Additionally, since ∇_**x**_ **· u** = 0, the last term in the integral above is also 0. Hence:

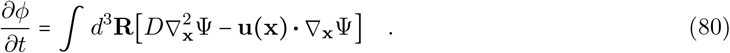

Finally, pulling the integral through the ∇_**x**_ and **u**, we have:

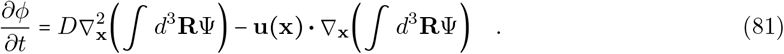

Finally, recognizing the definition of *ϕ* from Eq. (75), we arrive at:

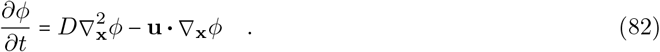

#### 1.8 Metrics for quantifying simulations

In order to quantitatively compare simulations as parameters are varied, we developed metrics to quantify different aspects of interest in each of the simulations. These are Basal-Apical Offset (BA Offset), Apical Shear, and Apical-Basal Shear Ratio (A:B Shear). Basal-Apical Offset is a representative measurement of the how displaced the basal surface is from the apical, and approximately measures the vertical “slantedness” of the cells. Apical Shear is a representative measurement of the amount of vortical flow at the apical surface. Finally, Apical-Basal Shear Ratio is a representative measurement of how similar vortical flow is at the apical versus basal surfaces.

First, the centroid of a cell is calculated as the centroid of the 24 points which make up the cell boundary. We then define measurements used in calculating these metrics:

- 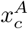 and 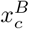 are the *x*-component of the displacement traveled by the centroid of the center cell (*c*) at the apical (*A*) and basal (*B*) surfaces.
- 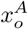 and 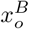 are the *x*-component of the displacement traveled by the northern-most, outer cell (*o*) at the apical (*A*) and basal (*B*) surfaces.

With these coordinates 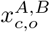, we calculate the metrics as follows.

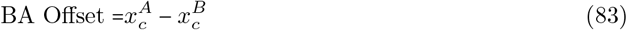

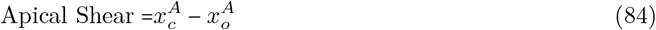

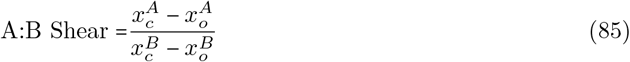

Based on measurements from fixed and live data, expected ranges for BA Offset is 0.25*r* to 1.2*r*, for Apical Shear is >2*r*, and for A:B Shear is 0.7 to 1.0, where *r* is the radius of a cell (*r* = 4 *µ*m for placode cells).

### 2 Simulating cell crawling with active forces

In order to simulate how cells move and monitor how cell shapes change, we track material points embedded in the fluid that represent the cell boundaries and move them via the velocity field. In the beginning of the simulation, we initiate these points in a pattern of vertical hexagonal prisms, each prism representing the initial shape of a cell. As the simulation progresses and these tracked points move, their new positions represent the deformed shape of a cell. We then utilize these points to specify the location of the crawling forces **F**^*t*^ applied to the cellular material. Because we are interested in testing the idea that inner cells provide the driving force, we only apply the crawling force to the central inner 19 cells.

The goal was to create an applied force profile for each cell that follows the shape and outline of that cell; see Fig. S5. Therefore, for each cell, the applied force profile was calculated based on the tracked points representing that cell’s boundaries at the basal surface; we represent these as 2D points since their *z* coordinate is simply 0. There are 24 of these points per cell: *P* = {**p**_1_, **p**_2_,…, **p**_24_}, where **p**_*i*_ = (*x*_*i*_, *y*_*i*_) . First, the lateral mid-point **p**_m_ is calculated from the average of the left-most and right-most points **p**_l_ and **p**_r_ in *P* as:

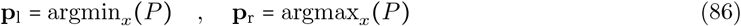

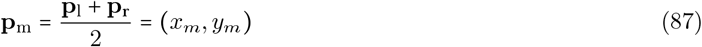

Here, the functions argmin_*x*_ (*P*) and argmax_*x*_ (*P*) simply return the point **p** ∈ *P* that has the minimum or maximum *x*-coordinate value in *P* . The position **p**_m_ allows us to then define the points which comprise the anterior *P*_*A*_ and posterior *P*_*P*_ regions of each cell.

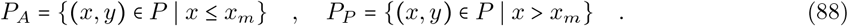

Following this, for each set of boundary points *P*_*A*_ and *P*_*P*_ representing the two halves of the cell, we compute the left- and rightmost, as well as the top- and bottom-most points in each of these regions:

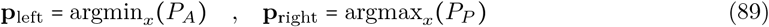

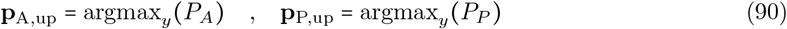

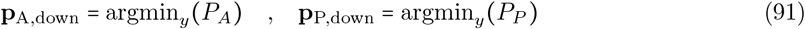

In order to specify the applied force to match the radius of the cell as the cell shape changes, we calculate the length *L* and width *W* of each half of the cell. The width of the anterior region *W*_*A*_ is defined as the distance between **p**_m_ and the left-most point of the anterior **p**_left_, while the width of the posterior *W*_*P*_ is defined as the distance between **p**_m_ and the rightmost point of the posterior **p**_right_. The lengths *L*_*A*_ and *L*_*P*_ are defined as distance between the top- and bottom-most points of the anterior and posterior regions:

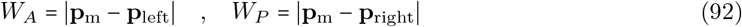

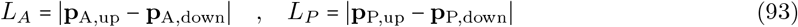

We compute angles of rotation *θ*_*A*_ and *θ*_*P*_ in the anterior and posterior regions so that if the cell outline appears rotated, then the force profile also becomes rotated to follow the shape of the cell. Let **p**_left_ = (*x*_left_, *y*_left_) and **p**_right_ = (*x*_right_, *y*_right_). We utilize the function atan2 (*y, x*), the 2-argument actangent function to compute:

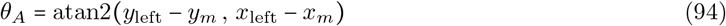

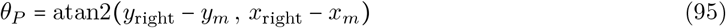

Note that if the cell is mostly unrotated, then *θ*_*A*_ ≈ *π* and *θ*_*P*_ ≈ 0. We apply this rotation to the coordinate system and positions within the cells by defining the rotation matrix **R**. We denote **x**^′*A*^ and **x**^′*P*^ as coordinates that have gone through rotations by *θ*_*A*_ and *θ*_*P*_, respectively:

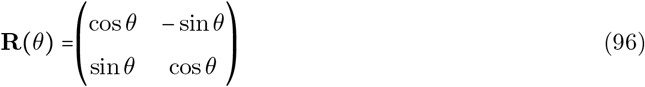

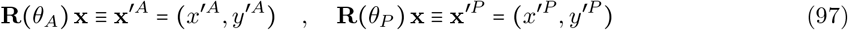

The forces are applied so as to follow the geometry of the cell given the lengths, widths, and rotation calculated above. Additionally, the duration of the force application is during a period *T* = 2. From times 0 to 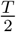, force is applied at the anterior half of the cell, and then from times 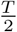 to *T*, force is applied in the posterior half of the cell. Finally, the magnitude of this force *F*_mag_ (*t*) is modulated over the period by

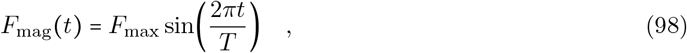

where *F*_max_ is the maximum force, a parameter in the simulation, and *t* is the time since the beginning of the force application. Finally, the shape of the force profile is that of an annulus, meaning that force is applied along the periphery of the cell; see Figure S5 and Eq. (99). The shape of the annulus is comprised of two gaussian functions, one that is wide and one that is narrow. The first pair of conditions in Eq. (99) is due to the *θ*_*A*_ ≈ *π* rotation where positions in the anterior of the cell are mapped to the positive *x*^′^ side of the coordinate system. These functions include a small correction factor *k* = 1.9, to ensure the perimeter of the cell is within the area of the force applied:

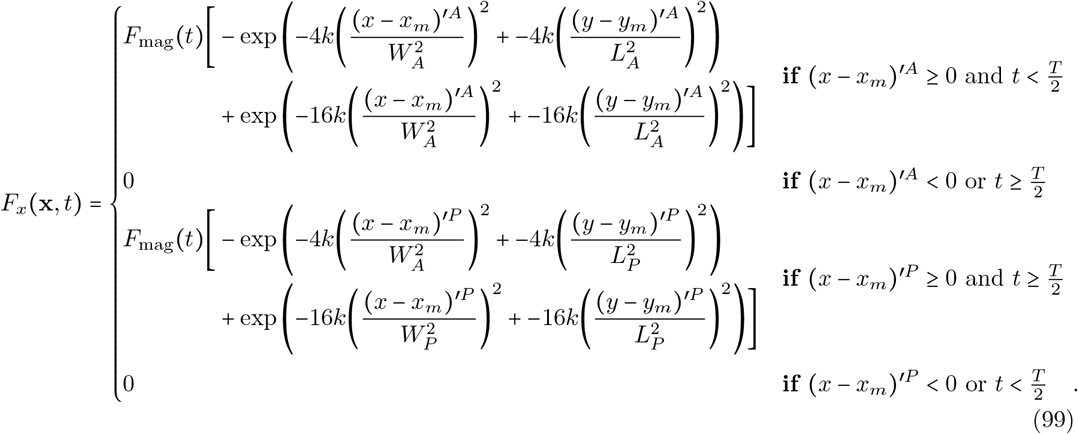

Finally, to allow for asynchronous crawling between the cells, each cell generates a random wait time from a random normal distribution *T*_wait_ ∼ *N* (*µ* = 0.5, *σ*^2^ = 0.25). Once the wait time has elapsed, the above described period of forces are applied. After a period of a crawl, the cell then generates a new wait time and the cycle repeats.

**Figure S5:**
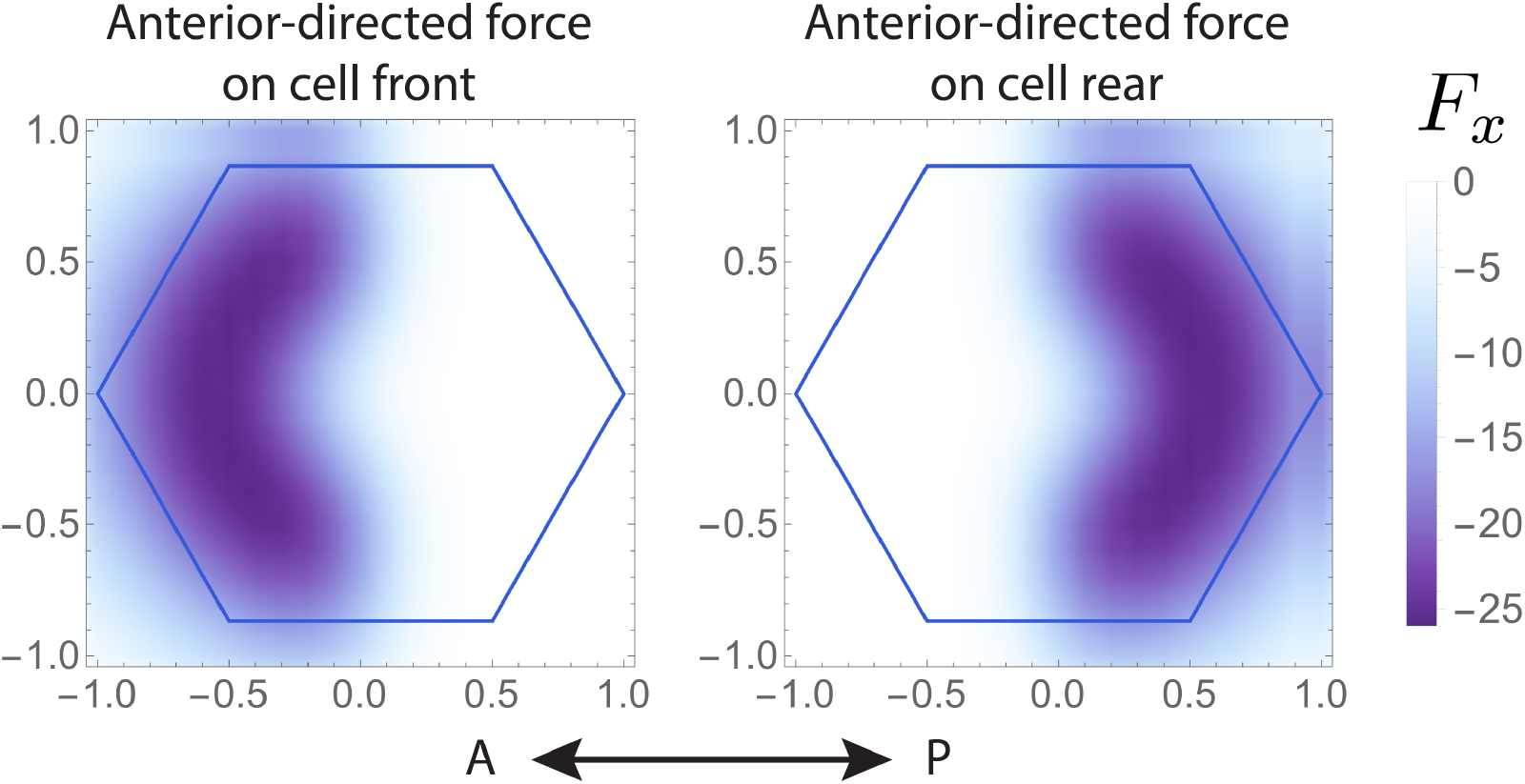
Active forces *F*_*x*_ applied in the negative *x* direction on each cell in the simulation. Anterior-directed forces on the cell front and rear are applied in turn, with forcing on the cell front occurring before the forcing on the cell rear.

